# Conformational buffering underlies functional selection in intrinsically disordered protein regions

**DOI:** 10.1101/2021.05.14.444182

**Authors:** Nicolas S. Gonzalez-Foutel, Wade M. Borcherds, Juliana Glavina, Susana Barrera-Vilarmau, Amin Sagar, Alejandro Estaña, Amelie Barozet, Gregorio Fernandez-Ballester, Clara Blanes-Mira, Ignacio E Sánchez, Gonzalo de Prat-Gay, Juan Cortés, Pau Bernadó, Rohit V. Pappu, Alex S. Holehouse, Gary W. Daughdrill, Lucía B. Chemes

## Abstract

Many disordered proteins conserve essential functions in the face of extensive sequence variation. This makes it challenging to identify the forces responsible for functional selection. Viruses are robust model systems to investigate functional selection and they take advantage of protein disorder to acquire novel traits. Here, we combine structural and computational biophysics with evolutionary analysis to determine the molecular basis for functional selection in the intrinsically disordered adenovirus early gene 1A (E1A) protein. E1A competes with host factors to bind the retinoblastoma (Rb) protein, triggering early S-phase entry and disrupting normal cellular proliferation. We show that the ability to outcompete host factors depends on the picomolar binding affinity of E1A for Rb, which is driven by two binding motifs tethered by a hypervariable disordered linker. Binding affinity is determined by the spatial dimensions of the linker, which constrain the relative position of the two binding motifs. Despite substantial sequence variation across evolution, the linker dimensions are finely optimized through compensatory changes in amino acid sequence and sequence length, leading to conserved linker dimensions and maximal affinity. We refer to the mechanism that conserves spatial dimensions despite large-scale variations in sequence as conformational buffering. Conformational buffering explains how variable disordered proteins encode functions and could be a general mechanism for functional selection within disordered protein regions.

## INTRODUCTION

Intrinsically disordered proteins and protein regions (IDRs) are abundant components of proteomes that play key roles in regulating cellular processes [^1,2^]. IDRs often bind their cellular partners via short linear motifs (SLiMs). These SLiMs represent conserved interaction modules that play key roles in cell signalling and can be identified from multiple sequence alignments [^3^]. In contrast, the contributions of seemingly unconcerned regions outside of SLiMs to IDR function are often less-well understood. Under the classical structure-function paradigm, the low sequence conservation and high frequency of insertions and deletions of IDRs [^4^] is indicative of weak evolutionary restraints, leading to the view that many IDRs might play the roles of passive “spacers”, stringing together ordered domains and disordered SLiMs. However, recent progress in the quantitative description of sequence-ensemble relationships in IDR conformations [^5^] indicates that specific features are required to support biological activity [^6,7,8,9^]. The observation that IDRs with vastly different sequence characteristics have conserved sequence-ensemble relationships (SERs) [^10^], suggests that SERs that determine function are under natural selection.

IDRs play major roles in viral adaptation by facilitating the acquisition of novel traits [^11,12,13^], making viral proteins ideal models to uncover the molecular mechanisms that shape functional evolution within IDRs. Tethering is a major function encoded by IDRs [^14^] that allows functional coupling between domains or SLiMs and regulates key processes including enzyme catalysis [^15^], transcriptional regulation [^16^] and liquid condensate formation [^17^]. The intrinsically disordered adenovirus early region 1A (E1A) protein is a multifunctional signalling hub that employs multiple SLiMs [^13,18,19^] tethered by disordered linkers to hijack cell signalling [^20^]. The subversion of cell cycle regulation by E1A involves crucial interactions with the retinoblastoma (Rb) tumour suppressor. Specifically, E1A uses two SLiMs [^21^] tethered by a disordered linker to bind to Rb and displace Rb-bound E2F transcription factors, triggering S-phase entry and viral genome replication at the low expression levels present hours after infection [^22^] (**Fig. 1a, b**). Additional binding motifs for the CREB binding protein (CBP) TAZ2 domain [^23^] and the BS69 transcriptional repressor MYND domain [^20^] (**Fig.1b**), mediate the formation of ternary complexes regulated by allostery [^24^]. Here we test the central hypothesis regarding conserved SERs driving functions in the disordered E1A linker, demonstrating that functionally equivalent IDRs that perform a tethering function can emerge despite dramatic changes in the linear sequence.

**FIGURE 1.**
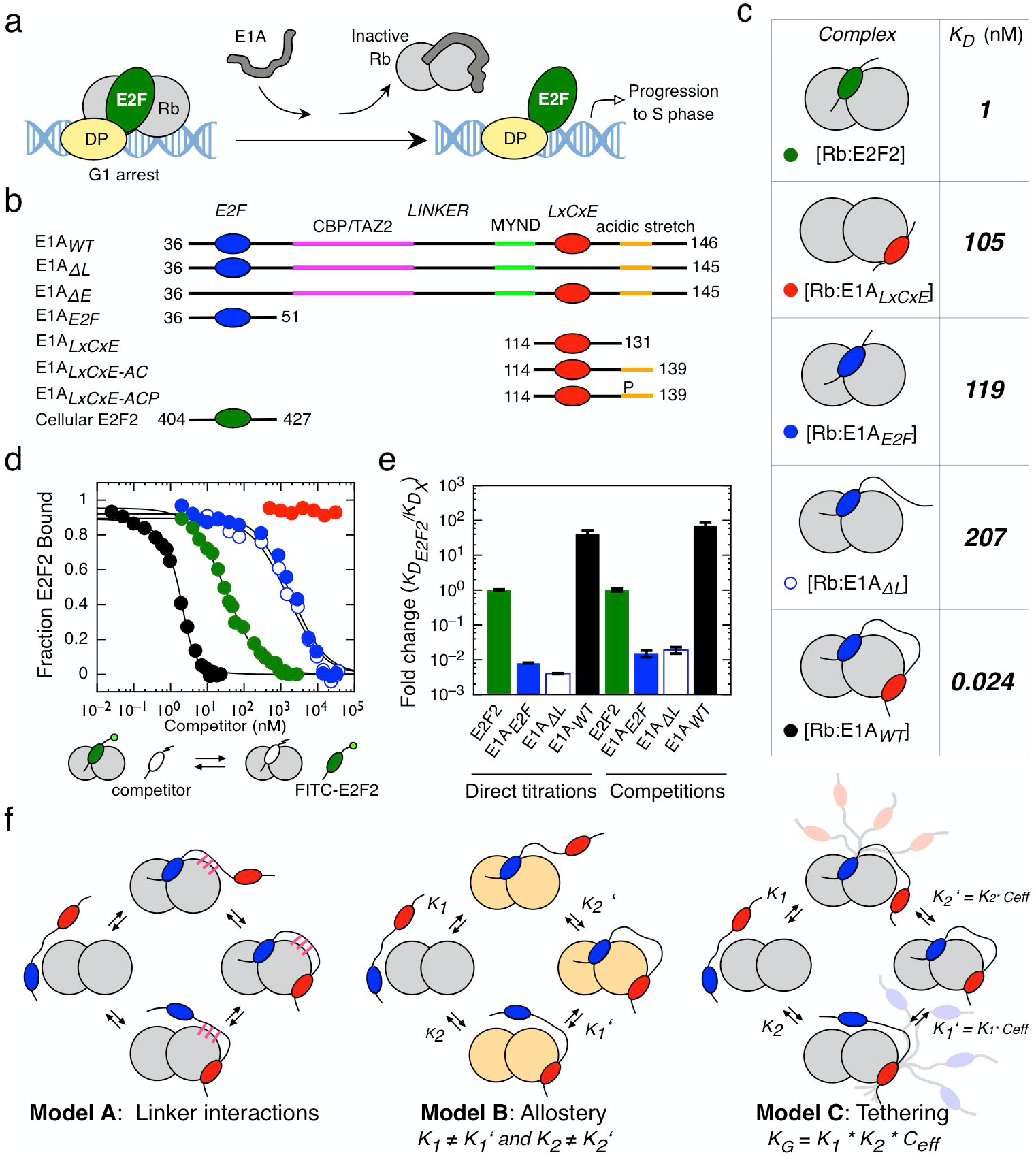
Tethering is required for high affinity Rb binding and E2F displacement by E1A. **a)** Model for disruption of the repressive Rb-E2F complex by E1A. **b)** Schematic representation of E1A and E2F2 constructs used in this study. Colour coding for the E2F, LxCxE, TAZ2 and MYND SLiMs, the acidic stretch and S132 phosphorylation are maintained throughout figures. **c)** Representative interactions tested using fluorescence spectroscopy (**Extended Data Fig. 2 and Extended Data Tables 1 and 2**). **d)** E2F competition titrations. Colour code is as in panel c. **e)** Comparison of the fold-change in binding affinity from direct titrations versus competition assays. Values higher than unity indicate an increase in binding affinity with respect to E2F2. **f)** Three models that account for affinity enhancement in the Motif-Linker-Motif E1A arrangement (See main text for details).

## RESULTS

### Tethering is required for high affinity Rb binding and E2F displacement

To uncover the molecular mechanisms underlying E2F displacement, we selected the minimal Rb binding region from the adenovirus serotype 5 (HAdV5) E1A protein. This region harbours the E1A_E2F_ and E1A_LxCxE_ SLiMs tethered by a 71-residue linker (E1A_*WT*_). We tested a series of E1A constructs comprising the individual motifs or fragments where the E2F (E1A_*ΔE*_) or LxCxE (E1A_*ΔL*_) motifs were mutated to poly-alanine (**Extended Data Fig. 1, Fig.1b**). For comparison we also tested the E2F SLIM (E2F2) taken from the host transcription factor E2F2 (**Fig. 1b**). Isothermal titration calorimetry (ITC) (**Extended Data Fig. 2 Extended Data Table 1**) and SEC-SLS experiments (**Extended Data Table 3**) confirmed that all E1A constructs bound to Rb with 1:1 stoichiometry. To quantify binding affinities, we performed fluorescence polarization measurements using FITC-labelled constructs (**Extended Data Fig. 3 Extended Data Tables 1 and 2**). While the host-derived E2F2 SLiM bound to Rb with high affinity (*K_D_* = 1 nM), the E1A_*E2F*_ SLIM had a *K_D_* = 119 nM, suggesting it would be a weak competitor of E2F2 (**Fig. 1c**). To our surprise, the motif-linker-motif arrangement (E1A_*WT*_) enabled binding to Rb with picomolar affinity (*K_D_* = 24 pM). This represents a 4000-fold enhancement compared to E1A_*E2F*_ and a 40-fold enhancement compared to E2F2.

It has been proposed that a minimal flexible linker or “spacer” is required to allow the simultaneous binding of both SLiMs that is required for the displacement of E2F [^25^]. To test the role of tethering in E2F displacement, we carried out competition assays. As expected, neither E1A_*LxCxE*_, nor E1A_*E2F*_ or E1A_*ΔL*_ were able to outcompete E2F from Rb (**Fig. 1d**). However, E1A_*WT*_ was a strong competitor, disrupting the [E2F2:Rb] complex at low nanomolar concentration (**Fig. 1d**). The agreement among ITC, direct titration and competition experiments confirmed that tethering was required for high affinity Rb binding and E2F displacement (**Fig. 1e, Extended Data Table 1**).

The striking affinity enhancement between the independent and linked SLiMs of E1A can be explained by three alternative models (**Fig. 1f**). In Model A, the E1A linker enhances affinity by establishing additional stabilizing interactions with the Rb domain. In Model B, a primary interaction by the E1A_*E2F*_ or E1A_*LxCxE*_ SLiMs induces an allosteric change in Rb that enables the complementary motif to bind with higher affinity. In Model C, once a primary interaction is established, the linker functions as an entropic tether that maximizes the effective concentration (*C_eff_*) of the second motif. We tested each of these models using a combination of biophysical measurements, molecular simulations, and evolutionary analysis.

### The disordered E1A linker does not make additional stabilizing interactions with Rb

We used NMR spectroscopy to determine the structural basis for the E1A_*WT*_ binding to Rb. For the E1A_*WT*_ in isolation, the transverse optimized relaxation (TROSY) spectrum of ^15^N-labeled E1A_*WT*_ revealed narrow chemical shift dispersion in the ^1^H-dimension. This is a characteristic signature of disordered regions and is consistent with previous work on E1A fragments (**Fig. 2a**) [^23^]. Further, the ^13^C_α_ secondary chemical shifts (C_α_Δδ) showed minimal deviation from random coil values obtained from disordered proteins (**Fig. 2b I**). Negative NHNOE values observed for E1A_*WT*_ indicated fast backbone dynamics (**Fig. 2b II**). Finally, sequence analysis also predicted that and E1A_*WT*_ is globally disordered (**Fig. 2b IV**). Taken together, these results indicate that the conformational ensemble of E1A_*WT*_ is characterized by high heterogeneity (disorder) and with fast interconversion between distinct conformations on the nanosecond to picosecond timescale (flexibility).

**FIGURE 2.**
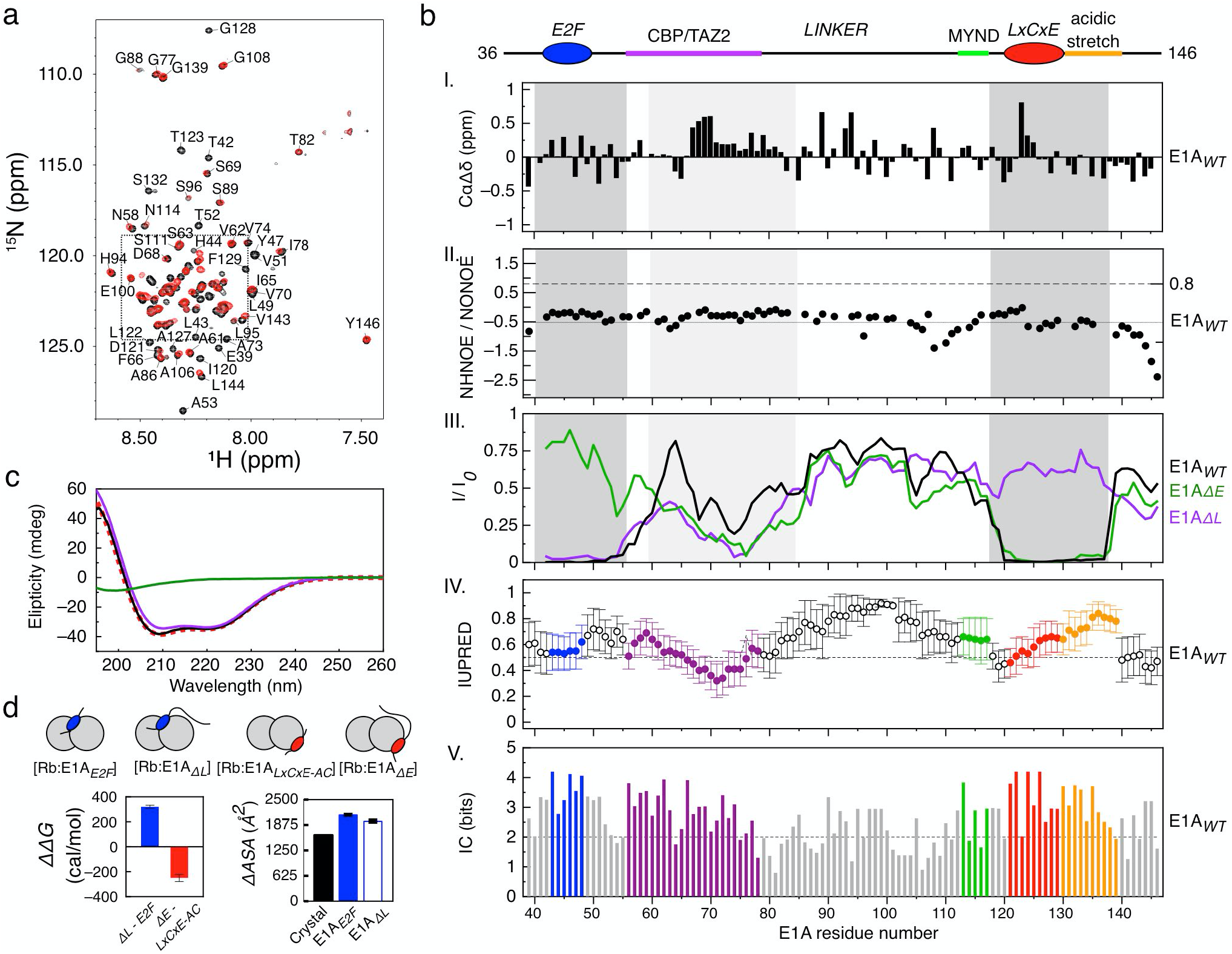
NMR and ITC analysis of the Rb-E1A interaction. **a)**^1^H-^15^N TROSY spectra of free 15N-E1A_*WT*_ (black) and ^15^N-E1A_*WT*_ bound to unlabelled Rb (red). ^15^N-E1A_*WT*_ peak assignments for the inset are shown in **Extended Data Fig. 4**. **b)**Ι. ^13^Cα secondary chemical shift (CαΔδ) of ^15^N-E1A_*WT*_. II. NHNOE/NONOE ratio for ^15^N-E1A_*WT*_. Dashed line: reference value for rigid backbone. IIΙ. Intensity ratio plots of bound state (*I*) with respect to the free state (*I_0_*) for E1A_*WT*_, E1A_*ΔL*_ and E1A_*ΔE*_. Dark grey: E2F/LxCxE SLiMs and flanking regions; Light grey: N-terminal linker region. **ΙV-V)** Disorder propensity and residue conservation (information content: IC) across 110 E1A sequences. **c)** Far-UV CD spectra for E1A_*WT*_ (green line), Rb (violet line), [Rb:E1A_*WT*_] complex (black line) and the arithmetic sum of the Rb and E1A_*WT*_ spectra (red dashed line). The latter CD spectra largely overlap. **d)** Changes in binding free energy (left) and *ΔASA* (right) for fragments containing or lacking the linker region.

Next, we examined the impact of binding of labelled E1A_*WT*_ to unlabelled Rb. The complex of E1A_*WT*_ and Rb is 54.6 KDa (**Extended Data Table 3**) and we expect to observe global chemical shift changes and line broadening for the regions of E1A_*WT*_ that interact with Rb. The TROSY spectrum of E1A_*WT*_ bound to Rb does not show any large chemical shift changes or widespread peak broadening for residues in the linker consistent with previous reports (**Fig. 2a and 2b III**) [^23^]. These results indicate that the linker region remains disordered and flexible when bound to Rb, an interpretation supported by the lack of changes in secondary structure upon binding (**Fig. 2c**). Based on our affinity data (**SI Text Section 1**) and previous reports [^26^], we anticipated that the region flanking the canonical E1A_*E2F*_ or E1A_*LxCxE*_ motifs contributes stabilizing interactions to the complex. In agreement with this expectation, the peaks corresponding to the E1A_*E2F*_ and E1A_*LxCxE*_ SLiMs (L43 to Y47 and L122 to E126) and their flanking residues (E39 to T52 and V119 to E135) disappeared upon binding, yielding near-zero I/I_0_ ratios (**Fig. 2b III**). This result is consistent with persistent intermolecular interactions, and independent binding of each motif to Rb (**SI Text Section 1**).

The N-terminal linker region encompassing the TAZ2 binding motif showed a decrease in peak intensities (**Fig. 2b III**) that could be due to stabilizing interactions with Rb [^23^]. However, an isolated E1A linker fragment (E1A_*60-83*_) showed no detectable association to Rb (**Extended Data Fig. 2i**) and E1A constructs including the linker region did not show higher binding affinities than isolated E1A motifs (**Fig. 2d, Extended Data Table 1**). Finally, the linker region did not contribute to the change in accessible surface area upon binding (**Fig. 2d, SI text Section 2, Extended Data Fig. 5h and Extended Data Table 4**). Collectively, these results demonstrate that the linker does not contribute to the thermodynamics of binding through coupled folding and binding or through discernible contacts with Rb. These results appear to rule out Model A (**Fig. 1f**).

### Affinity enhancement is not due to allosteric coupling between binding sites

To assess whether allosteric coupling between the E2F and LxCxE binding sites in Rb could explain the affinity enhancement (**Fig. 1f, Model B**), we performed ITC titrations where Rb was pre-saturated with E1A_*E2F*_ or E1A_*LxCxE*_ and titrated with the complementary motif (**Extended Data Fig. 5**). If a positive allosteric effect is at play, E1A_*LxCxE*_ should bind more tightly to Rb when E1A_*E2F*_ already bound and vice versa. This was measured as *ΔΔG* = *ΔG*_*PRE-SATURATED*_ − *ΔG_NON-SATURATED_*, where a negative value for *ΔΔG* indicates positive cooperativity. However, *ΔΔG* values for both motifs were lower than ± 0.25 kcal/mol (**Extended Data Table 5**). In E1A_*LxCxE*_ binding assays, pre-saturation with E1A_*ΔL*_ instead of E1A_*E2F*_ did not change the outcome, indicating that neither the motif nor the motif + linker arrangement behaved as allosteric effectors on the complementary site. Therefore, our results suggest that allosteric coupling is unlikely to be responsible for affinity enhancement. This rules out Model B (**Fig. 1f**) and points to tethering (**Fig. 1f, Model C**) as the mechanism underlying the ability of E1A to disrupt the E2F-Rb complex.

### The E1A linker acts as an entropic tether that optimizes the affinity of the E1A_LxCxE_ and E1A_E2F_ SLiMs for Rb

As suggested by early reports [^27^] tethering could allow docking through the E1A_*LXCXE*_ motif, increasing the effective concentration (*C_eff_*) of the E1A_*E2F*_ motif such that it efficiently outcompetes E2F (**Fig. 1f, Model C**). This form of “zeroth-order” cooperativity can be described using a simple Worm Like Chain (WLC) model that treats the linker as an entropic tether (**Fig. 3a,b**) [^28^]. According to this model, a short linker would be unable to straddle the distance between the two binding sites (**Fig. 3a,b l**), an optimal linker would maximize *C_eff_* (**Fig. 3a,b II**), and a longer than optimal linker would decrease *C_eff_* (**Fig. 3a,b III**). Application of the WLC model yields a predicted *C_eff_* value of 0.92 mM, with a near-optimal linker length (**Fig. 3b**). This is in close agreement with the estimate for *C_eff_* (0.52 ± 0.09 mM) obtained from the measured affinities, implicating model C (**Extended Data Table 1**).

**FIGURE 3:**
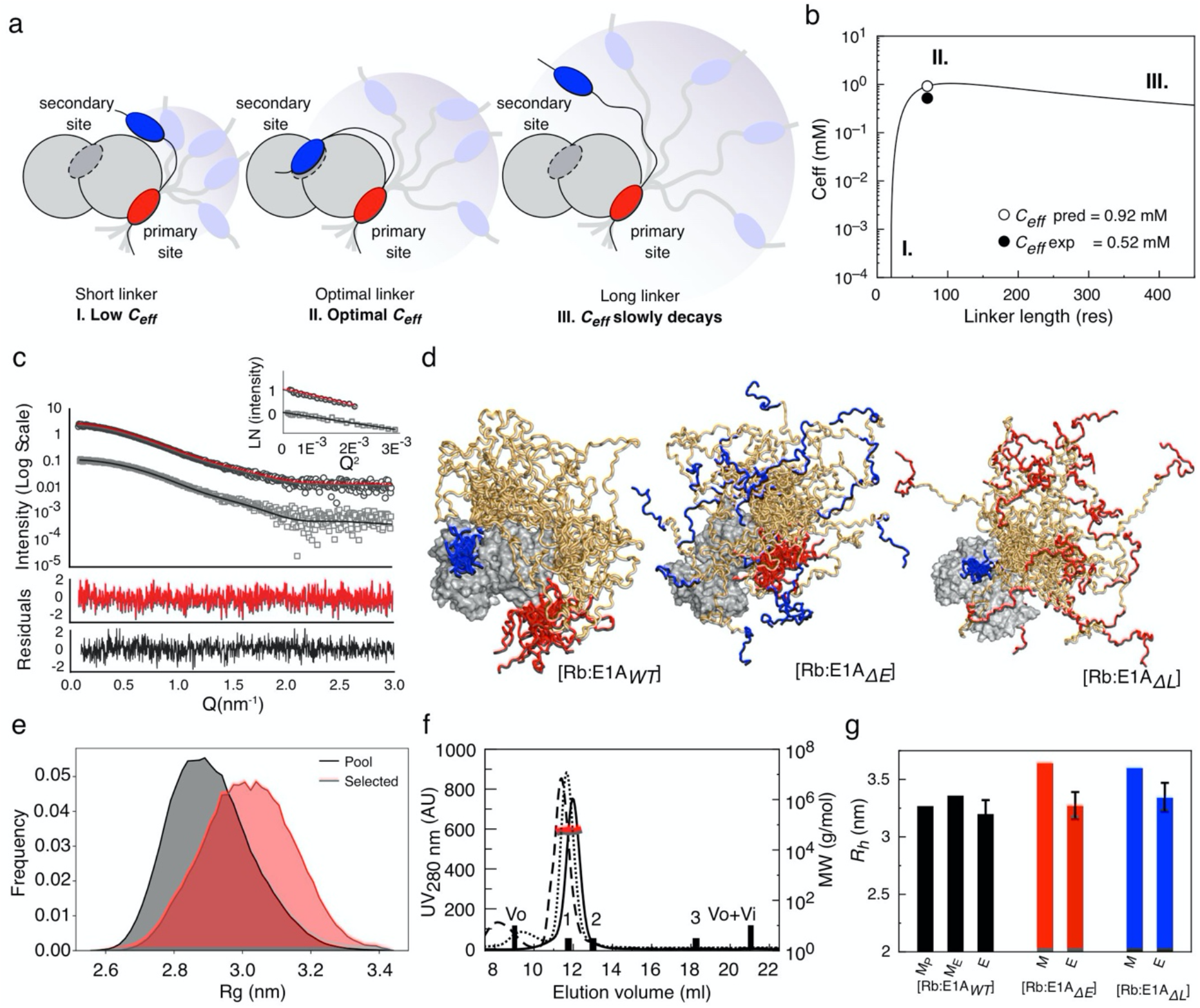
The E1A linker behaves as an entropic tether. **a)** Schematic representation of how *C_eff_* depends on linker length. **b)***C_eff_* curve from the WLC model. The scenarios depicted in a) are shown as regions (I, II, III). **c)** SAXS intensity profile of: Rb (grey squares) with best fit to the theoretical profile derived from the RbAB crystal structure (black line); and the [Rb:E1A_*WT*_] complex (black circles) with best fit from the EOM method (red line). Inset: Guinier plots for Rb and [Rb:E1A_*WT*_]. **d)** SAXS-selected [Rb:E1A_*WT*_] EOM ensemble (both motifs bound) and simulated ensembles for [Rb:E1A_*ΔE*_] and [Rb:E1A_*ΔL*_] (one motif bound). **e)** R_*g*_ distribution of the ensemble pool for [Rb:E1A_*WT*_] (black) and the EOM ensemble (red). The linker samples conformations more extended than the random-coil model of the pool. **f)** SEC-SLS of [Rb:E1A_*WT*_] (solid line), [Rb:E1A_*ΔE*_] (dotted line) and [Rb:E1A_*ΔL*_] (dashed line). Black bars: BSA 66 kDa (1), MBP 45 kDa (2) and Lysozyme 14.3 kDa (3). Black line: SEC profile, Red line: MW value (g/mol). **g)** Comparison between the hydrodynamic radius (R_*h*_) of modelled (M_P_ = pool, M_E_= EOM) and experimental (E) ensembles for [Rb:E1A_*WT*_] (black bars), [Rb:E1A_*ΔE*_] (red bars) and [Rb:E1A_*ΔL*_] (blue bars).

To assess the prediction that the E1A linker behaves as an entropic tether, we performed Small Angle X-ray Scattering (SAXS) experiments on Rb, *E1A_WT_*, and the [Rb:E1A_*WT*_] complex (**Fig. 3c, Extended Data Fig. 6**). The experimental SAXS profile of the RbAB domain could be fitted to the theoretical SAXS profile derived from its crystal structure (χ_i_^2^ = 1.3) and further refined (RMSD = 1.7 Å) using a SAXS-driven modelling approach (χ_i_^2^ = 0.82) (**Fig. 3c, Extended Data Fig. 6a**), indicating that Rb in solution retained its folded structure. Instead, the Kratky plots of *E1A_WT_* were characteristic of an IDP. Fitting of the SAXS profiles using the Ensemble Optimization Method (EOM) [^29^] indicated *E1A_WT_* was highly expanded (**Extended Data Fig. 6b**). To analyse the conformation of the linker in the [Rb:E1A_*WT*_] complex, we applied a sampling method [^30^] to generate a pool of 10250 realistic conformations [^31^] and computed theoretical SAXS profiles that were selected using EOM analysis. The SAXS profile of the complex was best described by sub-ensembles where the linker sampled expanded conformations (**Fig. 3c-e, Extended Data Fig. 6c**) with *R_h_* values (*R_h EOM_* = 3.36 nm) matching those obtained from SEC-SLS experiments (*R_h SEC_* = 3.20 ± 0.12 nm) (**Fig. 3f-g, Extended Data Fig. 6d and Extended Data Table 3**) and *R_g_*/*R_h_* ratios consistent with compaction upon bivalent tethering (**SI Text Section 3**). Collectively, the WLC modelling and the NMR, SAXS and SEC-SLS data support Model C, showing that the E1A linker behaves as an entropic tether that remains highly flexible and expanded to optimize the affinity of both binding motifs to Rb (**Fig. 1f**).

### Conformational buffering leads to conservation of the E1A linker dimensions

Inspection of selected linker sequences representative of *Mast adenoviruses* that infect a variety of mammalian hosts revealed significant variations in linker length and sequence composition (**Extended Data Fig. 7a**). While N- and C-terminal acidic extensions and an aromatic/hydrophobic TAZ2 binding region were highly conserved (**Fig. 2b V**), the linker lengths and compositions vary considerably within the central region enriched predominantly with polar, hydrophobic and proline residues. To gain insight into how these variations impacted linker conformations, we performed all atom simulations [^8^] and obtained conformational ensembles of 24 E1A linker sequences that exhaustively sample linker sizes ranging from 41 to 75 residues (**Source Data File 1**). Strikingly, the average end-to-end distance of these linkers remained roughly constant despite an almost doubling of the length (**Fig. 4a**), indicating the global linker dimensions were preserved through compensatory changes in sequence length and composition. We refer to this adaptive mechanism as *conformational buffering*. To uncover the sequence-encoded origins of conformational buffering we examined various statistical properties (**Extended Data Fig. 7b**). Net charge per residue (NCPR) was the strongest predictor of normalized end-to-end distance, with more expanded chains having a higher NCPR (**Fig. 4b**). Longer chains tend to have higher proline contents with fewer hydrophobic and charged residues (**Extended data fig. 7b, Fig. 4b, inset**). Our results suggest that conformational buffering is achieved through compensatory covariations in sequence that preserve the mean end-to-end distances across linker sequences. To test this hypothesis directly, we performed simulations for a collection of 140 random synthetic sequences of variable length that matched the amino acid composition of one of the shortest representative linkers (HF_HAdV40). In sharp contrast to natural sequences, the synthetic sequences showed a clear monotonic increase in end-to-end distance with chain length (**Extended Data Fig. 7c**). This result suggests that the sequence composition and patterning within natural E1A linkers are decidedly non-random, being tuned throughout evolution to ensure that changes in composition and chain length did not significantly alter the key physical properties of the linker. Our findings underscore the functional implications of preserving sequence-ensemble-relations (SERs), which in the case of E1A is achieved by preserving the dimensions of disordered linkers in order to enable the hijacking of the eukaryotic cell cycle by the virus.

**FIGURE 4.**
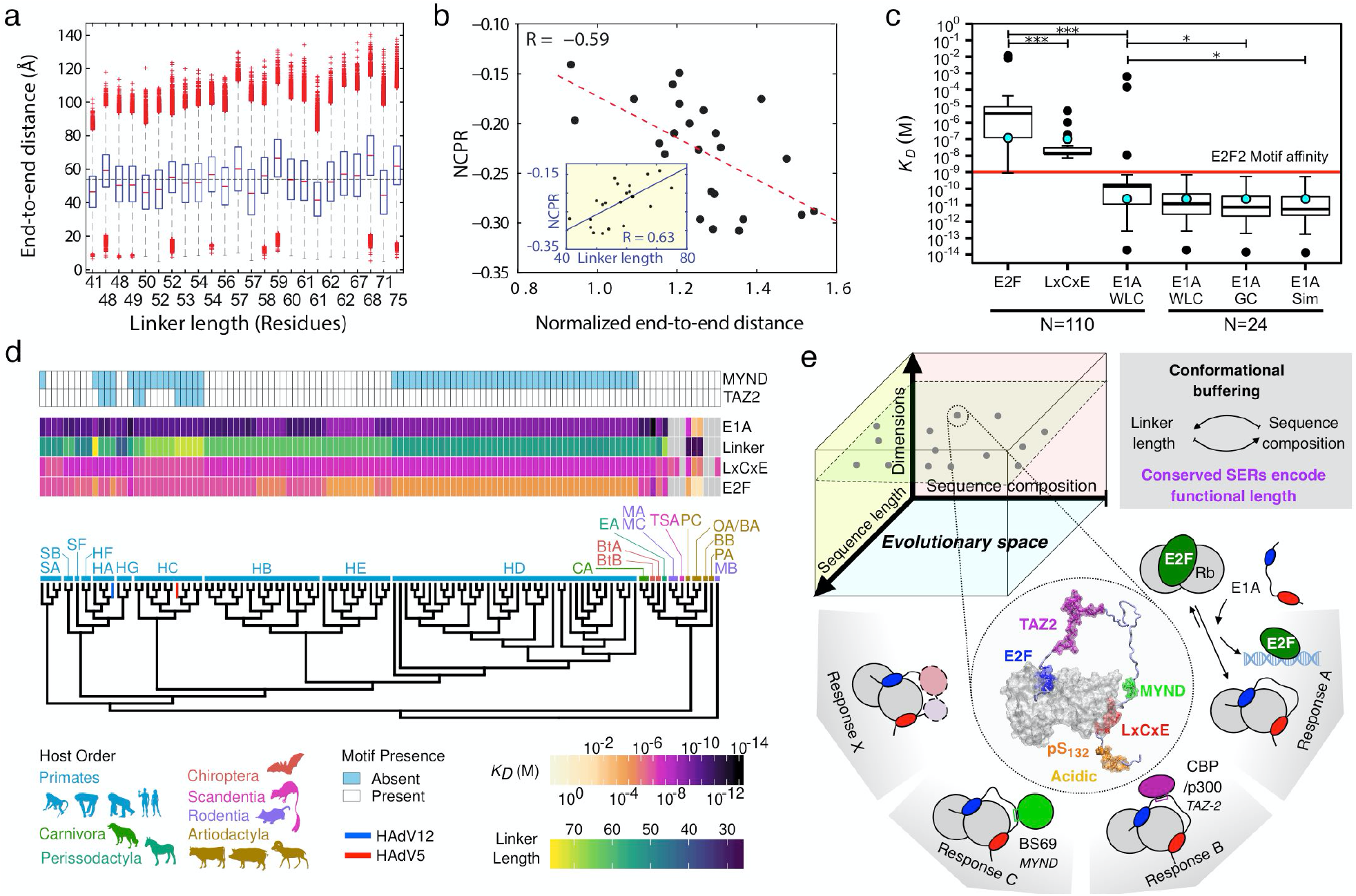
Evolutionary conservation of tethering. **a)** End-to-end distance calculated from all-atom simulations using a set of 24 E1A linker sequences. Horizontal dotted line: mean end-to-end distance (53.39 Å). **b)** Net-charge per residue (NCPR) as a function of normalized end-to-end distance. Inset: Net-charge per residue (NCPR) as a function of linker length. **c)***K_D_* values for the E1A_E2F_ and E1A_LXCXE_ SLiMs and E1A_*WT*_ for 110 sequences and 24 sequences used in simulations. Black dots: outliers, cyan dots: experimental value for HAdV5-E1A. *K*_*D,E1A*_ was calculated using different values for the *L_P_* parameter (**Extended Data Fig. 8 and Methods**), * *p*-value<0.05, *** *p*-value<0.001. Red line: affinity of cellular E2F2. **d)** Phylogenetic tree of *Mastadenovirus* E1A proteins with species denoted by two letter codes. The affinity of the E2F/LxCxE SLiMs and *E1A_WT_*, and linker length are indicated by Colour scales. Grey box: absent motif/linker. Light/blue box: present TAZ2/MYND SLiMs. **e)** Upper: E1A sequences evolved a multiplicity of solutions in the sequence space to achieve conserved SERs through conformational buffering. Lower: The model represents one pose of the conformational [Rb:E1A_WT_] ensemble with E2F/LxCxE SLiMs bound to Rb. The evolvable E1A interaction platform performs highly conserved functions (E2F activation) while allowing adaptive changes in functionality.

### Evolutionary conservation of E1A tethering

If tethering is a key functional feature, then the *C_eff_* and global Rb binding affinity should be under selection. Accordingly, the expectation is that these parameters should be conserved across E1A evolution. In agreement with this expectation, the predicted *C_eff_* values for 110 E1A linkers covering the *Mastadenovirus* E1A phylogeny were distributed close to the maximum of the *C_eff_* function (**Extended Data Fig. 8a, c**). Next, we calculated the global Rb binding affinity of all E1A proteins (*K_D,E1A_*) using the predicted *C_eff_* values together with individual motif affinities predicted using energy scoring matrices (**Extended Data Fig. 8d**). Strikingly, the median values of 110 E1A proteins followed the trend observed in HAdV5E1A: while the E1A_*E2F*_ motif bound Rb with weaker affinity (*K_D,E2F_* = 3.9 μM) than the cellular E2F2 motif (*K_D,E2F2_* = 1 nM) (**Fig.4c**, red line), the global binding affinity (*K*_*D,E1A*_ = 150 pM) was higher than E2F2 suggesting that the hijack mechanism is conserved across E1A proteins (**Fig. 4c**).

The compensatory changes uncovered by our all-atom simulations suggested that linker-specific C_*eff*_ may confer tighter binding affinities when compared to a naïve WLC model in which chain length is the sole determinant of C_*eff*_. The relevant parameter is the apparent chain stiffness or persistence length (*L_p_*) (**Extended Data Fig. 8b**). While *L_p_* is kept constant in the naïve WLC model, in reality the apparent stiffness varies between sequences in a composition-dependent manner [^28^]. In order to explore how conformational buffering affected the C_*eff*_ enforced by the linkers, we calculated sequence-specific *L_p_* values (*L_p_*Sim) from the all-atom simulations (**See Methods**). The median *L_p_*Sim value (6.7 Å) was higher than the naïve expectation (*L_p_* = 3Å) (**Extended Data Fig. 8e**). Thus, E1A linkers were globally expanded [^32^], and approached an optimal linker length (**Extended Data Fig. 8f,g**). While the naïve WLC model predicts *C_eff_* to increase from 0.4 mM to 1 mM from the shortest to the longest linker, chain-specific *L_p_*Sim values led to all linkers displaying a *C_eff_* of 1 mM (**Extended Data Fig. 8h,i**). This produced a 3-fold enhancement in the global binding affinity for short linkers (**Extended Data Fig. 8j**) and improved the median *K*_*D,E1A* Sim_ value to 5.9 pM (**Fig. 4c**) independent of variations in sequence length (**Extended Data Fig. 8k**). Accordingly, we propose that the *functional length* of the linker [^33^] referred to as a joint contribution of sequence length, amino acid composition, and sequence patterning as determinants of end-to-end distances – is not random. Instead, the functional length is conserved through conformational buffering to enable maximal affinity enhancement.

Phylogenetic analysis further revealed that tethering was strongly conserved within primate-infecting *Mast adenoviruses* and host orders Carnivora, Chiropteran and Perissodactyla (**Fig. 4d**). In contrast, tethering was weakened within more divergent orders (PC, OA, BA, **Extended Data Fig. 7a** and outlier points in **Fig. 4c**) due to the presence of short linkers coupled to low affinity or absent Rb binding motifs (**Fig. 4d**). This suggests that the motif-linker-motif arrangement is under co-evolutionary selection, such that either the linker and the motifs are jointly optimized, or neither are under selective pressure, presumably leading to a loss of the displacement mechanism.

## DISCUSSION

Here, we demonstrate how E1A hijacks the eukaryotic cell cycle using two SLiMs tethered by an optimal entropic linker [^34,35^]. The proposed docking and displacement mechanism appears to be maintained by natural selection through conformational buffering, a mechanism that promotes robust encoding of core functions while supporting the extensive variation of sequence features that would be required for viral adaptation. Our work challenges the view that IDRs evolve with few restrictions, demonstrating that distinct features, specifically sequence ensemble relationships (SERs) of IDR dimensions are likely to be under selective pressure that can be masked by sequence variation and naïve sequence alignments.

For the E1A tethers, the main feature driving conformational buffering is net charge per residue (NCPR), in agreement with previous findings that charge valence and patterning are major determinants of IDR dimensions in natural [^33,36,37,38^] and synthetic [^37,39,40^] sequences. Additional linker features might help maintain specific conformations: the acidic extensions may generate electrostatic repulsion that ensures the linker stays extended, exposing the associated TAZ2 and MYND motifs, while favoring local solvation that prevents linker-Rb interactions. Previous studies have shown that proline can contribute more or less [^40^] strongly to IDR compaction. Changes in proline content were not a major determinant of E1A linker expansion, but instead appear to reflect an orthogonal selection feature such as the inhibition of helical secondary structure (**Extended Data Fig. 7**). This underscores the reality that multiple mechanisms might contribute to fine-tune the dimensions of E1A linkers. Our results suggest that WLC models coupled with sequence-based estimations of persistence length could be used to create more realistic representations of multivalent interactions in systems biology toolboxes [^41^].

We uncover how conformational buffering maintains the linker functional length through compensatory changes in sequence length, composition, and patterning that give rise to a diverse set of functionally equivalent IDRs (**Fig. 4e**, upper). This sequence variability could reflect adaptive changes following host switch events [^42^] that support rewiring of the E1A interactome through the gain and loss of SLiMs [^19,43^] or enhance viral immune evasion by fine tuning interactions with immune suppressors that map onto the region of interest [^44^] (**Fig. 4e,** lower). The robust encoding of functions provided by conformational buffering could be widespread among IDRs, underlying gene silencing [^6^], kinase inhibitor function [^7^], transcriptional control [^16,36^] and phase separation [^8^].

## Data deposition

The EOM ensembles have been deposited in the Protein Ensemble Database (proteinensemble.org) with codes PED00174 (E1A_WT_:Rb) and PED00175 (E1A_WT_)

## Acknowledgements

This work was supported by Agencia Nacional de Promoción Científica y Tecnológica (ANPCyT) Grants PICT 2013-1895 and 2017-1924 (to LBC), 2012-2550 and PICT 2015-1213 to IES and 2016-4605 (to GPG), the US National Institutes of Health (grants GM115556 and CA141244 to GWD, grant 5R01NS056114 to RVP), FLDOH grant 20B17 (to GWD), and the US National Science Foundation (grant MCB-1614766 to RVP). GWD and LBC were supported by a travel award from the USF Nexus Initiative and a Creative Scholarship Grant from the USF College of Arts and Sciences. PB was supported by the Labex EpiGenMed, an «Investissements d’avenir» program (ANR-10-LABX-12-01) (to PB). The CBS is a member of France-BioImaging (FBI) and the French Infrastructure for Integrated Structural Biology (FRISBI), two national infrastructures supported by the French National Research Agency (ANR-10-INBS-04-01 and ANR-10-INBS-05, respectively). This work was supported by the Spanish Ministerio de Ciencia y Universidades (MICYU-FEDER, RTI2018-097189-C2-1 to GFB) NGF was supported by a doctoral fellowship from Consejo Nacional de Investigaciones Científicas y Técnicas (CONICET, Argentina) and a scholarship from Fulbright Visiting Scholar Program. NGF and JG are CONICET postdoctoral fellows, and LBC, IES and GdPG are CONICET researchers. SBV was supported by fellowships from Ministerio de Ciencia e Innovación, España #BES-2013-063991 and #EEBB-I-16-11670. ASH is supported by the Longer Life Foundation: A RGA/Washington University Collaboration. This work benefited from the HPC resources of the CALMIP supercomputing center under the allocation 2016-P16032 and the Cluster of Scientific Computing (http://ccc.umh.es/) of the Miguel Hernández University (UMH). We thank Kathryn Perez at the Protein Expression and Purification Core Facility at EMBL (Heidelberg) for critical help with ITC experiments.

## Author Contribution Statement

LBC, GWD, ASH and RVP designed research and conceived the study. NSGF and WMB produced reagents. NSGF and WMB performed FP, ITC and NMR experiments and WB, NGF, GWD and LBC analysed data. JG conducted bioinformatic analyses of E1A variants. AS and PB performed and analysed SAXS experiments. AE, AB and JC produced and analysed E1A conformational ensembles. SBV and ASH performed and analysed all atom simulations of E1A linkers. GFB, CBM and IES computed and analysed FOLDX matrices. NSGF, JG, AS and ASH produced figures. LBC, GWD, PB, JC, GdPG, IES, ASH and RVP Supervised research. LBC, NSGF, JG, RVP, ASH and GWD wrote the paper with critical feedback from all authors.

## METHODS

### Protein purification and peptide synthesis and labeling

#### Protein expression and purification

The human Retinoblastoma protein (Uniprot ID: P06400) AB domain with a stabilizing loop deletion (372-787Δ582-642), named Rb, was recombinantly expressed from a pRSET-A vector in *E. coli* Bl21(DE3) following a previously described protocol [^1^]. The adenovirus serotype 5 (HAdV5) Early 1A protein fragment (36-146) (Uniprot ID: P03255), named E1A_*WT*_, was subcloned into *Bam*HI/*Hin*dIII sites of a modified pMalC vector (NewEnglandBioLabs, Hitchin, UK). E1A_*ΔE*_ (43-LHELY-47Δ43-AAAA-46) and E1A_ΔL_ (122-LTCHE-126Δ122-AAAA-125) variants were obtained by site-directed mutagenesis of the wild type vector. E1A proteins were expressed as MBP fusion products in *E. coli* BL21(DE3). Unlabelled and single (^15^N) and double (^15^N/^13^C) labelled samples were obtained from 2TY medium and M9-minimal medium supplemented with ^15^NH_4_Cl and ^13^C-glucose respectively. Cultures were induced with 0.8 mM IPTG at 0.7 *OD*_600_ and grown at 37 °C overnight in 2TY medium or for 5 h after induction in M9-minimal medium. Harvested cells were lysed by sonication and proteins isolated performing amylose affinity chromatography of the soluble fraction, followed by Q-HyperD Ion exchange and size exclusion (Superdex 75) chromatography. The MBP tag was cleaved with Thrombin (Sigma-Aldrich, USA) at 0.4 unit per mg of protein. Protein purity (> 90%) and conformation were assessed by SDS-PAGE, SEC-SLS and circular dichroism analysis (**Extended Data Fig. 1**).

#### Peptide synthesis

Peptides corresponding to individual E1A or E2F2 binding motifs were synthesized by FMoc chemistry at >95% purity (GenScript, USA) and quantified by Absorbance at 280 nm or by quantitation of peptide bonds at 220 nm in HCl -when Tryptophan or Tyrosine residues were absent. The peptide sequences are:

E1A_*E2F*_ 36-SHFEPPTLHELYDLDV-51

E1A_*LxCxE*_ 116-VPEVIDLTCHEAGFPP-131

E1A_*LxCxE-AC*_ 116-VPEVIDLTCHEAGFPPSDDEDEEG-139

E1A_*LxCxE-ACP*_ 116-VPEVIDLTCHEAGFPP*pS* DDEDEEG-139

Human E2F2 404-SPSLDQDDYLWGLEAGEGISDLFD-427

#### FITC labeling

Proteins and peptides were labelled at their N-terminus with Fluorescein 5-Isothiocyanate (FITC, Sigma), purified and quantified following a described protocol [^45^]. F/P (FITC/Protein) ratio was above 0.8 in all cases.

### Circular Dichroism

Far-UV CD spectra were measured on a Jasco J-810 (Jasco, Japan) spectropolarimeter equipped with a Peltier thermostat using 0.1 or 0.2 cm path-length quartz cuvettes (Hellma, USA). Five CD scans were averaged from 195 to 200 nm at 100nm/min scan speed, and buffer spectra were subtracted from all measurements. All spectra were measured in 10mM Sodium Phosphate buffer pH 7.0 and 2mM DTT at 20 ± 1 °C and 5 μM protein concentration.

### Size Exclusion Chromatography, Hydrodynamic radii calculations and Light Scattering Experiments

Analytical size exclusion chromatography (SEC) was performed on a Superdex 75 column (GE Healthcare) calibrated with globular standards: BSA (66 kDa), MBP (45 kDa) and Lisozyme (14.3 kDa). All runs were performed by injecting 100 μl protein sample (E1A_*WT*_ and E1A_*ΔL*_ at 270 μM and E1A_*ΔE*_ at 540 μM) in 20 mM Sodium Phosphate buffer pH 7.0, 200 mM NaCl, 2 mM DTT. For each protein or complex a partition coefficient (K_av_) was calculated and apparent molecular weights were interpolated from the –logMW vs K_av_ calibration curve. Experimental hydrodynamic radii (*R_h_*) were calculated following empirical formulations developed by Uversky and col. [^46^]:

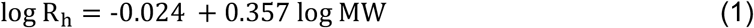

Where *MW* is the apparent molecular weight derived from SEC experiments. The predicted *R_h_* for E1A_*WT*_ was calculated following the formulation developed by Marsh and Forman-Kay [^3^].

The exponent *ν* was calculated from *R_h_* = *R_o_*·*N^ν^* using the experimental *R_h_* values, with *R_o_* = 2.49 nm for E1A_*WT*_ and *R_o_* = 4.92 nm for Rb, following [^32^]. For E1A_*WT*_, ν was calculated from R_*g*_ = R_*0*_·*N*^ν^ using *R_g_* obtained from SAXS measurements and R_*0*_ = 2.1 nm, following [^47^]. In both cases, *N* is the number of residues in the chain (**Extended Data Table 3**).

Static Light Scattering (SLS) coupled to SEC was carried out to determine the average molecular weight of individual protein peaks and the stoichiometry of [Rb:E1A] complexes using a PD2010 detector (Precision Detectors Inc, China) coupled in tandem to an HPLC system and an LKB 2142 differential refractometer. The 90° light scattering (LS) and refractive index (RI) signals of the eluting material were analysed with Discovery32 software (Precision Detectors).

Dynamic Light Scattering (DLS) was used to measure the hydrodynamic size distribution of E1A, using a Wyatt Dynapro Spectrometer (Wyatt Technologies, USA). Data was fitted using Dynamics 6.1 software. All measurements were performed in 20 mM Sodium Phosphate buffer pH 7.0, 200 mM NaCl, 1 mM DTT at 2 mg/ml. Samples were filtered by 0.22 μM filters (Millipore) and placed into a 96 Well glass bottom black plate (In Vitro Scientific P96-1.5H-N) covered by a high performance cover glass (0.17+/-0.005mm) before measurements were taken.

### Fluorescence Spectroscopy Experiments

Measurements were performed in a Jasco FP-6200 (Nikota, Japan) spectropolarimeter assembled in L geometry coupled to a Peltier thermostat. Excitation and emission wavelengths were 495 nm and 520 nm respectively, with a 4 nm bandwidth. All measurements were performed in 20 mM Sodium Phosphate buffer pH 7.0, 200 mM NaCl, 2 mM DTT and 0.1% Tween-20 at 20 ± 1 °C.

For direct titrations, a fixed concentration of FITC-labelled protein/peptide was titrated with increasing amounts of Rb until saturation was reached. Maximal dilution was 20% and samples equilibrated for 2 min ensuring steady state. Titrations performed at concentrations 10 times higher than the equilibrium dissociation constant (*K_D_*) allowed estimation of the stoichiometry of each reaction. Binding titrations performed at sub-stoichiometric concentrations allowed an estimation of *K_D_*, by fitting the titration curves to a bimolecular association model:

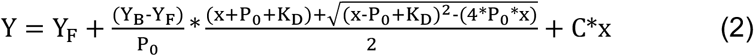

Where Y is the measured anisotropy signal, Y_F_ and Y_B_ are the free and bound labelled peptide signals, P_0_ is the total labelled peptide concentration, x is Rb concentration, and *K_D_* is the equilibrium dissociation constant in Molar units. The [C * *x*] linear term accounts for slight bleaching or aggregation. Data was fitted using the Profit 7.0 software (Quantumsoft, Switzerland), yielding a value for each parameter and its corresponding standard deviation. Titrations for each complex were performed in triplicate at least at three different concentrations of FITC-labelled sample, and parameters were obtained from fitting individual titrations or by global fitting of the *K_D_* parameter using normalized titration curves at different concentrations, obtaining an excellent agreement between individual and global fits (**Extended Data Table 2**).

Competition experiments were carried out by titrating the pre-assembled complex [Rb:FITC-E2F2] (1:1 molar ratio, 5 nM) with increasing amounts of unlabelled competitors and following the decrease in the anisotropy signal until the value corresponding to free FITC-E2F2 was reached. IC50 values were estimated directly from the curves as the concentration where the competitor produced a decrease in 50% of the maximal anisotropy value. *K_D_* values were calculated by fitting the data considering the binding equilibrium of the labelled peptide and the unlabelled competitors according to [^48^], obtaining *K_D(comp)_* values that differed only slightly (2 to 4-fold) from those obtained from direct titrations. *K_D_* and *K_D(comp)_* values also displayed similar fold changes in binding affinity relative to E2F2 within each method (**Extended Data Table 1**). The agreement between the *K_D_* values obtained from fluorescence and ITC titrations (**Extended Data Table 1**) confirmed that FITC moiety did not cause significant changes in Rb binding affinity.

### ITC Experiments

#### Direct titrations

ITC experiments were performed on MicroCal VP-ITC and MicroCal PEAQ-ITC equipment (Malvern Panalytical) in 20 mM Sodium Phosphate pH 7.0, 200 mM NaCl, 5mM 2-mercapto ethanol at 20.0 ± 0.1 °C, unless stated otherwise. Prior to titrations, cell and titrating samples were co-dialyzed in the aforementioned buffer for 48 h at 4 ± 1 °C and then de-gassed. Measurements performed in the MicroCal VP-ITC used 28 10-μl injections at a flow rate of 0.5 μl/s and those performed in the MicroCal PEAQ-ITC used 13 3-μl injections. The concentration range of cell and titrating samples are detailed in **Extended Data Figs. 2 and 5**. Data were analysed using the Origin software.

#### Allosteric coupling experiments

First, a pre-assembled [Rb:E1A_*LxCxE*_] complex (1:1 molar ratio, 30 μM) was titrated with E1A_*E2F*_ or E1A_*ΔL*_ to assess whether binding of the LxCxE motif modified the binding affinity for the E2F site. Conversely, pre-assembled [Rb: E1A_*E2F*_] or [Rb: E1A_*ΔL*_] complexes were titrated with E1A_*LxCxE*_ to assess whether binding of the E2F motif modified the binding affinity for the LxCxE site (**Extended Data Table 5**).

#### Calculation of ΔCp and ΔASA parameters from ITC data

A series of titrations were carried out at different temperatures (10.0, 15.0, 20.0 and 30.0 ± 0.1 °C) and the change in binding heat capacity (*ΔCp*) was obtained from the slope of the linear regression analysis of the plot of *ΔH* vs temperature (**Extended Data Fig. 5**). The changes in accessible surface area and the number of residues involved in the interaction were estimated by solving semi-empirical equations from protein unfolding studies applied to protein-ligand binding [^49^](**Extended Data Table 4**).

### NMR Experiments

NMR experiments were carried out using a Varian VNMRS 800 MHz spectrometer equipped with triple resonance pulse field *Z*-axis gradient cold probe. A series of two-dimensional sensitivity-enhanced ^1^H–^15^N HSQC and three-dimensional HNCACB, HNCO and CBCA(CO)NH experiments [^50,51^] were performed for backbone resonance assignments on uniformly ^13^C–^15^N-labelled samples of E1A_*WT*_, E1A_*ΔE*_ and E1A_*ΔL*_ at 700 μM, 975 μM and 850 μM respectively. All measurements were performed in 10 % D2O, 20 mM Sodium Phosphate pH 7.0, 200 mM NaCl, 2 mM DTT at 25 °C. The HSQC used 9689.9 Hz and 1024 increments for the t1 dimension and 2106.4 Hz with 128 increments for the t2. The HNCACB used 9689.9, 14075.1, and 2106.4 Hz, with 1024, 128, and 32 increments for the t1, t2, and t3 dimensions, respectively. The HNCO used 9689.9, 2010.4 Hz, and 2106.4 Hz with 1024, 64, and 32 increments for the t1, t2, and t3 dimensions, respectively. The CBCA(CO)NH used 9689.9, 14072.6, and 2106.4 Hz, with 1024, 128, and 32 increments for the t1, t2, and t3 dimensions, respectively. For E1A_*WT*_ 88% of non-proline backbone ^1^H and ^15^N nuclei, 75% of ^13^C’ nuclei and 90% of ^13^C_α_ and ^13^C_β_ of E1A nuclei were assigned. For E1A_*ΔE*_ and E1A_*ΔL*_ 85% of non-proline backbone ^1^H and ^15^N nuclei, 72% of ^13^C’ nuclei and 87% of ^13^C_α_ and ^13^C_β_ E1A nuclei were assigned.

NMRPipe and NMRViewJ software packages were used to process and analyse all the NMR spectra [^52^]. Residue-specific random coil chemical shifts were generated for the three sequences using the neighbor-corrected IDP chemical shift library [^53^]. Secondary chemical shifts (Δ δ), were calculated by subtracting random coil chemical shifts from the experimentally obtained chemical shifts.

Two-dimensional ^1^H–^15^N TROSY experiments were performed on single ^15^N-labelled samples of free E1A_*WT*_, E1A_*ΔE*_ and E1A_*ΔL*_ and on each E1A protein bound stoichiometrically to Rb (1:1 molar ratio) at 525 μM (E1A_*WT*_), 300 μM (E1A_*ΔE*_) and 315 μM (E1A_*ΔL*_). The ratio between the peak intensity in the bound state (I) and the peak intensity in the free state (I_0_) was calculated, allowing interacting residues to be determined together with additional data.

### WLC modelling

#### The worm like chain (WLC) model

A worm like chain (WLC) model [^54^] was used to describe the end-to-end probability density distribution function of the E1A linker and estimate the effective concentration term (C_*eff*_) used in the tethering model (**Figure 1, Model C and Figure 3**). In this model, the disordered linker behaves as a random polymer chain whose dimensions depend on the persistence length (*L_P_*), which represents the chain stiffness, or the length it takes for the chain motions to become uncorrelated and on the contour length (*L_C_*), which is the total length of the chain. For long peptides, *L_P_* assumes a standard value of 3Å and *L_C_* is the product of the number of linker residues times the average unit size of one amino acid (3.8 Å) [^28^]. Under this model, the probability density function p(*r*) is defined by:

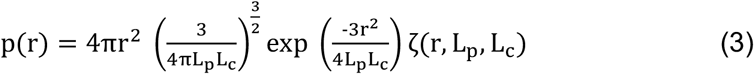

Where p(*r*) is a function of distance *r* and depends on *L_P_* and *L_C_*. The last term in the equation is expanded in [^54^,^28^]. The end-to-end probability density function can be related to the effective concentration in the bound state when the linker is restrained to a fixed distance between binding sites, *r*_o_ [^54^]. In this case, the effective concentration *C_eff_* is defined by:

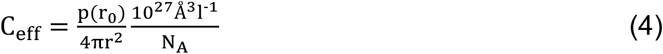

Where *N_A_* is Avogadro’s number and (*r_0_*) is the distance separating the binding sites obtained from the X-ray structure of the complex (49 Å calculated from PDB: 2R7G [^25^] and 1GUX [^55^]). Multiplying Eq. (4) by 10^3^ yields *C_eff_* in millimolar units.

#### Calculation of predicted C_eff_ value for the E1A_WT_:Rb interaction

The *C_eff_* value predicted from the WLC model was obtained by applying Eq. [4] with the designated *L_p_* parameter (standard model *L_P_* = 3Å and *b* = 3.8Å), using a linker length of 71 residues for HAdV5 E1A. The separation between binding sites, *r_0_*, was 49 Å (from PDB:1GUX and PDB:2R7G). The WLC model was also used to estimate the C_*eff*_ values of a collection of 110 natural linker sequences of different length changing the length value for each linker and keeping other parameters constant.

#### Calculation of sequence-dependent L_p_ values

In order to represent the extension of E1A linkers taking into account sequence composition, we derived the persistence length from all atom simulations (*L_p Sim_*). *L_p Sim_* was calculated from the average end-to-end distance of each simulated ensemble using the equation <r^2^>= 2\**L_p*_L_c_*, where *L_c_* = N*b and b takes the value 3.8 Å. This equation is an approximation for the value of <r^2^> for a worm like chain in the case where the contour length of the chain is much larger than its persistence length (*L_c_* >> *L_p_*) (**Source Data File 2**) [^28^]. New *C_eff_* values were derived using the same parameters described above, but replacing the standard *L_p_* value by the *L_p Sim_* value.

#### Calculation of experimental C_eff_ value for the E1A_WT_:Rb interaction

We calculated the experimental *C_eff_* value from Model C: *K_G_* = *K_1_*\**K_2_*\*C_*eff*_ (**Figure 1f**) where *K_G_*, *K_1_* and *K_2_* were the equilibrium association constants (*K* =1/*K_D_*) for binding of the motif-linker-motif construct E1A_*WT*_ (*K_G_*) or the individual motifs E1A_*E2F*_ (*K_1_*) and E1A_*LxCxE*_ (*K_2_*) to Rb (**Extended Data Table 1**). The condition *K_1_* = *K_1_*’ and *K_2_* = *K_2_*’ (no allosteric coupling between sites) was met, as experimentally proven (**Extended Data Fig. 5 and Extended Data Table 5**).

### Molecular modelling of Rb:E1A conformational ensembles

Conformations of E1A_*WT*_ bound to Rb were modelled using an extended version of a recently proposed method to generate realistic conformational ensembles of IDPs [^56^]. This method exploits local, sequence-dependent structural information encoded in a database of three-residue fragments and builds conformations incrementally sampling dihedral angles values from the database, while avoiding steric clashes. In order to model the double-bound [Rb:E1A_*WT*_] complex, the E2F and LxCxE motifs were considered to be static, preserving the conformations extracted from experimentally determined structures (2R7G and 1GUX). The 71-residue fragment between these two motifs was considered as a long protein loop that adapts its conformation in order to maintain the two ends rigidly positioned. Conformational sampling considering such loop-closure constraints was performed using a robotics-inspired algorithm [^31^] adapted to use dihedral angle values from the aforementioned database. For each feasible conformation of the central fragment, geometrically compatible conformations of the short N- and C-terminal tails were sampled using the basic strategy explained in [^56^]. For singly bound models, only one of the two motifs were considered to be statically bound to Rb and the other motif behaved as the flexible linker.

### SAXS Experiments

SAXS experiments were carried out at the European Molecular Biology Laboratory beamline P12 of DORIS and PETRAIII storage rings respectively, using the X-ray wavelengths of 1.24 Å and a sample-to-detector distance of 3.0 m. The scattering profiles measured covered a momentum transfer range of 0.0026 < *s* < 0.73 Å^−1^. SAXS data were measured for Rb, E1A_*WT*_ and [Rb:E1A_*WT*_] complex at 10° C. Concentrations used for E1A_*WT*_ were 7.0, 5.6 and 4.2 mg/ml, for Rb were 4.0, 2.0, 1.0 mg/ml, and for and [Rb:E1A_*WT*_] were 2.7, 1.4, and 0.7 mg/ml, in 20 mM Sodium Phosphate pH 7.0, 200 mM NaCl, 1mM DTT. The scattering patterns of the buffer solution were recorded before and after the measurement of each sample. Multiple repetitive measurements were performed to detect and correct for radiation damage. Final curves at each concentration were derived after the averaged buffer scattering patterns were subtracted from the protein sample patterns. No sign of aggregation was observed in any of the curves. Final SAXS profiles for the systems were obtained by merging curves for the lowest and highest concentrations to correct small attractive interparticle effects observed. Raw data manipulation was performed using standard protocols with the suite of programs ATSAS [^57^]. The forward scattering intensity, *I*(0), and the radius of gyration, *R_g_*, were evaluated using Guinier’s approximation [^58^], assuming that at very small angles (*s* < 1.3/*R_g_*, the intensity can be well represented as *I*(*s*) = *I*(0) exp(−(*sR_g_*)^2^/3)). The *P(r)* distribution functions were calculated by indirect Fourier Transform using GNOM [^59^] applying a momentum transfer range of 0.01 < *s* < 0.33 Å^−1^ and 0.013 < *s* < 0.27 Å^−1^ for Rb and [Rb:E1A], respectively. For E1A_*WT*_ a SEC-SAXS experiment was also performed to obtain the SAXS profile from a highly monodisperse sample. This profile overlaid perfectly with the E1A_*WT*_ merged curve from the three batch experiments, discarding aggregation problems.

The fitting of the crystallographic structure of Rb (PDB: 3POM [^60^]) to the experimental SAXS curve was performed with FOXS [^61^,^62^]. An optimal fit (χ^2^=0.86) was obtained after modelling the missing parts (loops, N- and C-termini) and a subsequent refinement with the program AllosMod-FoXS [^63^]. SAXS data measured for [Rb:E1A] were analysed with the Ensemble Optimization Method (EOM) [^29^,^64^]. Briefly, theoretical SAXS profiles of the 10250 structures of the complex were computed with CRYSOL [^65^]. 200 different sub-ensembles of 20 or 50 conformations collectively describing the experimental curve were collected with EOM and analysed in terms of *R_g_* distributions. The experimental SAXS data of [Rb:E1A_*WT*_] complex is compatible with three distinct scenarios: a 100% doubly-bound ensemble where the linker is highly expanded, a 100% singly-bound ensemble where the linker is highly compact and thirdly, an ensemble with a combination of 76% doubly bound and :24% singly-bound species, which resulted from the linear combination of a curve representing the ensemble average of all singly- and all doubly-bound conformations. However, thermodynamic (*K_D_* for *E1A_WT_*) data strongly argue against the last two scenarios as it indicates an extremely low expected population of the singly-bound forms at any concentration of the complex used in the SAXS experiments.

### Hydrodynamic radii for generated conformations

Hydrodynamic radii were calculated using the program HYDROPRO (version 10) [^66,67^]. HYDROPRO was run on 1000 models selected by EOM for the doubly-bound conformations and 1000 randomly selected conformations of N- and C-terminal bound conformations. The calculations were done at temperatures of 20 and 25 °C with corresponding solvent viscosities of 0.01 and 0.009 poise, respectively. The values of atomic element radius (AER), Molecular Weight, Partial Specific Volume and Solvent Density were set to 2.9 Å, 54590 Da, 0.702 cm^3^/g and 1.0 g/cm^3^, respectively.

### All-atom simulations of E1A Linker sequences

All-atom simulations were run using the CAMPARI simulation engine (V2) and ABSINTH implicit solvent model [^68,69^]. All simulations were run at 320 K; while this is a slightly elevated temperature compared to the experimental temperature, none of the terms the Hamiltonian lacks temperature dependence such that this slightly high temperature serves to improve sampling quality in a uniform way across all simulations. This approach has been leveraged to great effect in previous studies and is especially convenient in the case of simulating many different sequences that span a range of sequence properties and lengths [^7^]. A collection of Monte Carlo moves was used to fully sample conformational space as previously described [^70,71,36^].

For all simulations of natural sequences, 15 independent simulations were run per sequence for a total of 90K conformations per sequence across 27 different sequences (405 independent simulations, 5.25 ×10^8^ Monte Carlo steps per sequence). Simulations were performed in 15 mM NaCl in a simulation droplet size sufficiently large for each sequence, calibrated in a length dependent manner. Simulations were analysed using CTraj (http://ctraj.com), an analysis suite built on the MDTraj package [^72^]. Sequence analysis was performed using the local CIDER software package [^73^] with all parameters reported in (**Source Data File 1**). Normalized end-to-end distance was calculated as the absolute end-to-end distance divided by the end-to-end distance expected for an equivalently long Gaussian chain.

### Length titration Simulations

The linker from HF_HAdV40 was used to determine the overall amino acid composition and generate random sequences across a range of lengths that recapitulated this composition. Specifically, for each length (45, 50, 55, 60, 65, 70, 75) twenty random sequences were generated for a total of 140 randomly generated sequences. Each sequence was simulated under equivalent simulation conditions for 35 × 10^9^ simulation steps, with the goal of elucidating the general relationship between sequence length and end-to-end distance for an arbitrary sequence of the composition associated with HF_HAdV40. The mean end-to-end distance for the collection of sequences at a given length was determined, such that the mean value is a double average over both conformational space and sequence space.

### Calculation of global binding affinity of natural Adenovirus E1A sequences

#### Dataset

A previously reported alignment of 116 Mastadenovirus E1A sequences [^43^] was used to identify the E2F and LxCxE motifs as described [^43^], collecting 110 sequences in which both motifs were present (**Source Data File 4**). For all sequences, the length of the linker region between both motifs was recorded. Individual motif binding affinities, *C_eff_* values and E1A global affinity (*K_D,E1A_*) were calculated as explained below (**Source Data File 2**).

#### Calculation of E1A binding affinity

The global binding affinity (*K_D,E1A_*) was calculated from the predicted *C_eff_* values and the predicted binding affinities of each motif (*K_D,E2F_* and *K_D,LxCxE_*) from Model C (**Figure 1f**) as *K_D,E1A_* = *K_D,E2F_*\**K_D,LxCxE_*\**C_eff_*^−1^. Each term was calculated as follows:

#### Motif binding affinity prediction

To estimate the binding affinity of individual E2F and LxCxE motifs present in each sequence, FoldX 5.0 [^74^] was used to build substitution matrices for all 20 amino acids at each position (**Source Data File 2**). Briefly, given a structural complex the FoldX algorithm assesses the change in binding free energy produced by mutating each position of the motif for each one of the 20 amino acids. For the E2F matrix, the structure of the HAdV5 E1A_E2F_ motif in complex with Rb (PDB: 2R7G) was used as input. For the LxCxE matrix, the structure used as input was a model of the HAdV5 E1A_*LxCxE*_ motif in complex with Rb (**Source Data File 3**), built using FlexPepDock [^75^] and the structure of the HPV E7 LxCxE motif bound to Rb (PDB: 1GUX). The total change in binding free energy with respect to the wild type sequence (*ΔΔG*_FoldX_) was calculated by adding up the free energy terms for each residue at each matrix position (**Source Data File 2**). The predicted equilibrium dissociation constant of the E2F and LxCxE motifs for each sequence (*K_D SEQ_*) was calculated as:

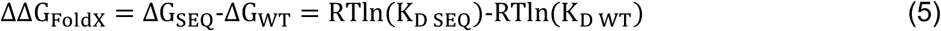

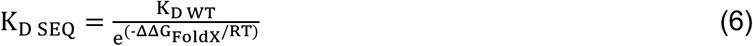

Where *ΔΔG*_FoldX_ is the total predicted change in binding energy calculated using FoldX, *RT* is 0.582 kcal mol^−1^, *K_D WT_* is the experimentally measured binding affinity of the sequence (HAdV5 E1A) present in the model structure (*K_D,E2F_* and *K_D,LxCxE_* measured in this work, **Extended Data Table 1**).

#### C_eff_ prediction

The *C_eff_* value was calculated for a collection of 110 natural E1A linkers as described in the “****WLC modelling****” section using Equation (4) and *L_p_* = 3 Å (*L_p_* WLC). For the subset of 24 natural E1A linkers used in all-atom simulations, we additionally calculated *L_p_* values obtained from the best fit of the data to a Gaussian Chain model (*L_p_* GC) and sequence-specific *L_p_* values from all atom simulations (*L_p_* Sim) as described in “****WLC modelling****”. All data are reported in (**Source Data File 2**).

#### Statistical analysis

We used bootstrapping [^76^] to generate 99% confidence intervals (CI) for *K_D,_*_E2F_, *K_D,_*_LxCxE_ and *K_D,_*_E1A_ average values, and compared the lower and upper end points against the value of *K_D,_*_E2F2_ (1 10^−9^ M). The lower bound of the 99% CI for *K_D_*_,E2F_ and *K_D_*_,LxCxE_ is higher than *K_D_*_,E2F2_ and the upper bound of the 99% CI for all *K_D_*_,E1A_ are lower than *K_D_*_,E2F2_. We also used permutation tests [^76^] to assess the null hypothesis that the *C_eff_*, *L_p_* and average *K_D_* average values did not differ between all pairs of groups. In order to control for the false discovery rate, the p-values were corrected using the Benjamini-Hochberg [^77^] correction for multiple comparisons.

#### Calculations of disorder propensity and conservation

All calculations were performed on the dataset from Source data file 4, using the methods described in [^43^]. For disorder propensity we recorded the mean IUPRED value ± SD per position and for residue conservation we recorded the information content (IC) per position.

## Notes

### Competing Interest Statement

The authors have declared no competing interest.

## REFERENCES

1. Wright PE, Dyson HJ. Intrinsically unstructured proteins: re-assessing the protein structure-function paradigm. J Mol Biol. 1999;293(2):321–331. doi:10.1006/jmbi.1999.3110

2. van der Lee R, Buljan M, Lang B, et al. Classification of intrinsically disordered regions and proteins. Chem Rev. 2014;114(13):6589–6631. doi:10.1021/cr400525m

3. Tompa P, Davey NE, Gibson TJ, Babu MM. A million peptide motifs for the molecular biologist. Mol Cell. 2014;55(2):161–169. doi:10.1016/j.molcel.2014.05.032

4. Brown CJ, Johnson AK, Dunker AK, Daughdrill GW. Evolution and disorder. Curr Opin Struct Biol. 2011;21(3):441–446. doi:10.1016/j.sbi.2011.02.005

5. Das RK, Ruff KM, Pappu R V. Relating sequence encoded information to form and function of intrinsically disordered proteins. Curr Opin Struct Biol. 2015;32:102–112. doi:10.1016/j.sbi.2015.03.008

6. Beh LY, Colwell LJ, Francis NJ. A core subunit of Polycomb repressive complex 1 is broadly conserved in function but not primary sequence. Proc Natl Acad Sci U S A. 2012;109(18):E1063–71. doi:10.1073/pnas.1118678109

7. Das RK, Huang Y, Phillips AH, Kriwacki RW, Pappu R V. Cryptic sequence features within the disordered protein p27Kip1 regulate cell cycle signalling. Proc Natl Acad Sci U S A. 2016;113(20):5616–5621. doi:10.1073/pnas.1516277113

8. Martin EW, Holehouse AS, Peran I, et al. Valence and patterning of aromatic residues determine the phase behavior of prion-like domains. Science. 2020;367(6478):694–699. doi:10.1126/science.aaw8653

9. Daughdrill GW, Narayanaswami P, Gilmore SH, Belczyk A, Brown CJ. Dynamic behavior of an intrinsically unstructured linker domain is conserved in the face of negligible amino acid sequence conservation. J Mol Evol. 2007;65(3):277–288. doi:10.1007/s00239-007-9011-2

10. Zarin T, Strome B, Nguyen Ba AN, Alberti S, Forman-Kay JD, Moses AM. Proteome-wide signatures of function in highly diverged intrinsically disordered regions. Elife. 2019;8. doi:10.7554/eLife.46883

11. Tokuriki N, Oldfield CJ, Uversky VN, Berezovsky IN, Tawfik DS. Do viral proteins possess unique biophysical features? Trends Biochem Sci. 2009;34(2):53–59. doi:10.1016/j.tibs.2008.10.009

12. Gitlin L, Hagai T, LaBarbera A, Solovey M, Andino R. Rapid evolution of virus sequences in intrinsically disordered protein regions. PLoS Pathog. 2014;10(12):e1004529. doi:10.1371/journal.ppat.1004529

13. Hagai T, Azia A, Babu MM, Andino R. Use of host-like peptide motifs in viral proteins is a prevalent strategy in host-virus interactions. Cell Rep. 2014;7(5):1729–1739. doi:10.1016/j.celrep.2014.04.052

14. Huang Q, Li M, Lai L, Liu Z. Allostery of multidomain proteins with disordered linkers. Curr Opin Struct Biol. 2020;62:175–182. doi:10.1016/j.sbi.2020.01.017

15. Dyla M, Kjaergaard M. Intrinsically disordered linkers control tethered kinases via effective concentration. Proc Natl Acad Sci U S A. 2020;117(35):21413–21419. doi:10.1073/pnas.2006382117

16. Brodsky S, Jana T, Mittelman K, et al. Intrinsically Disordered Regions Direct Transcription Factor In Vivo Binding Specificity. Mol Cell. 2020;79(3):459–471.e4. doi:10.1016/j.molcel.2020.05.032

17. Harmon TS, Holehouse AS, Rosen MK, Pappu R V. Intrinsically disordered linkers determine the interplay between phase separation and gelation in multivalent proteins. Elife. 2017;6. doi:10.7554/eLife.30294

18. Davey NE, Trave G, Gibson TJ. How viruses hijack cell regulation. Trends Biochem Sci. 2011;36(3):159–169. doi:10.1016/j.tibs.2010.10.002

19. Chemes LB, de Prat-Gay G, Sanchez IE. Convergent evolution and mimicry of protein linear motifs in host-pathogen interactions. Curr Opin Struct Biol. 2015;32:91–101. doi:10.1016/j.sbi.2015.03.004

20. King CR, Zhang A, Tessier TM, Gameiro SF, Mymryk JS. Hacking the Cell: Network Intrusion and Exploitation by Adenovirus E1A. MBio. 2018;9(3). doi:10.1128/mBio.00390-18

21. Dyson N, Guida P, McCall C, Harlow E. Adenovirus E1A makes two distinct contacts with the retinoblastoma protein. J Virol. 1992;66(7):4606–4611. http://www.ncbi.nlm.nih.gov/pubmed/1534854

22. Crisostomo L, Soriano AM, Mendez M, Graves D, Pelka P. Temporal dynamics of adenovirus 5 gene expression in normal human cells. PLoS One. 2019;14(1):e0211192. doi:10.1371/journal.pone.0211192

23. Ferreon JC, Martinez-Yamout MA, Dyson HJ, Wright PE. Structural basis for subversion of cellular control mechanisms by the adenoviral E1A oncoprotein. Proc Natl Acad Sci U S A. 2009;106(32):13260–13265. doi:10.1073/pnas.0906770106

24. Ferreon AC, Ferreon JC, Wright PE, Deniz AA. Modulation of allostery by protein intrinsic disorder. Nature. 2013;498(7454):390–394. doi:10.1038/nature12294

25. Liu X, Marmorstein R. Structure of the retinoblastoma protein bound to adenovirus E1A reveals the molecular basis for viral oncoprotein inactivation of a tumour suppressor. Genes Dev. 2007;21(21):2711–2716. doi:10.1101/gad.1590607

26. Palopoli N, Gonzalez Foutel NS, Gibson TJ, Chemes LB. Short linear motif core and flanking regions modulate retinoblastoma protein binding affinity and specificity. Protein Eng Des Sel. 2018;31(3):69–77. doi:10.1093/protein/gzx068

27. Fattaey AR, Harlow E, Helin K. Independent regions of adenovirus E1A are required for binding to and dissociation of E2F-protein complexes. Mol Cell Biol. 1993;13(12):7267–7277. http://www.ncbi.nlm.nih.gov/pubmed/8246949

28. Zhou HX. Polymer models of protein stability, folding, and interactions. Biochemistry. 2004;43(8):2141–2154. doi:10.1021/bi036269n

29. Bernado P, Mylonas E, Petoukhov M V, Blackledge M, Svergun DI. Structural characterization of flexible proteins using small-angle X-ray scattering. J Am Chem Soc. 2007;129(17):5656–5664. doi:10.1021/ja069124n

30. Estaña A, Sibille N, Delaforge E, Vaisset M, Cortés J, Bernadó P. Realistic Ensemble Models of Intrinsically Disordered Proteins Using a Structure-Encoding Coil Database. Structure. 2019;27(2):381–391.e2. doi:10.1016/j.str.2018.10.016

31. Cortes J, Simeon T, Remaud-Simeon M, Tran V. Geometric algorithms for the conformational analysis of long protein loops. J Comput Chem. 2004;25(7):956–967. doi:10.1002/jcc.20021

32. Marsh JA, Forman-Kay JD. Sequence determinants of compaction in intrinsically disordered proteins. Biophys J. 2010;98(10):2383–2390. doi:10.1016/j.bpj.2010.02.006

33. Cohan MC, Eddelbuettel AMP, Levin PA, Pappu R V. Dissecting the Functional Contributions of the Intrinsically Disordered C-terminal Tail of Bacillus subtilis FtsZ. J Mol Biol. 2020;432(10):3205–3221. doi:10.1016/j.jmb.2020.03.008

34. Borcherds W, Becker A, Chen L, Chen J, Chemes LB, Daughdrill GW. Optimal Affinity Enhancement by a Conserved Flexible Linker Controls p53 Mimicry in MdmX. Biophys J. 2017;112(10):2038–2042. doi:10.1016/j.bpj.2017.04.017

35. Bertagna A, Toptygin D, Brand L, Barrick D. The effects of conformational heterogeneity on the binding of the Notch intracellular domain to effector proteins: a case of biologically tuned disorder. Biochem Soc Trans. 2008;36(Pt 2):157–166. doi:10.1042/BST0360157

36. Sherry KP, Das RK, Pappu R V, Barrick D. Control of transcriptional activity by design of charge patterning in the intrinsically disordered RAM region of the Notch receptor. Proc Natl Acad Sci U S A. 2017;114(44):E9243–E9252. doi:10.1073/pnas.1706083114

37. Das RK, Pappu R V. Conformations of intrinsically disordered proteins are influenced by linear sequence distributions of oppositely charged residues. Proc Natl Acad Sci U S A. 2013;110(33):13392–13397. doi:10.1073/pnas.1304749110

38. Müller-Späth S, Soranno A, Hirschfeld V, et al. From the Cover: Charge interactions can dominate the dimensions of intrinsically disordered proteins. Proc Natl Acad Sci U S A. 2010;107(33):14609–14614. doi:10.1073/pnas.1001743107

39. Mao AH, Crick SL, Vitalis A, Chicoine CL, Pappu R V. Net charge per residue modulates conformational ensembles of intrinsically disordered proteins. Proc Natl Acad Sci U S A. 2010;107(18):8183–8188. doi:10.1073/pnas.0911107107

40. Sorensen CS, Kjaergaard M. Effective concentrations enforced by intrinsically disordered linkers are governed by polymer physics. Proc Natl Acad Sci U S A. 2019;116(46):23124–23131. doi:10.1073/pnas.1904813116

41. Bashor CJ, Horwitz AA, Peisajovich SG, Lim WA. Rewiring cells: synthetic biology as a tool to interrogate the organizational principles of living systems. Annu Rev Biophys. 2010;39:515–537. doi:10.1146/annurev.biophys.050708.133652

42. Hoppe E, Pauly M, Gillespie TR, et al. Multiple Cross-Species Transmission Events of Human Adenoviruses (HAdV) during Hominine Evolution. Mol Biol Evol. 2015;32(8):2072–2084. doi:10.1093/molbev/msv090

43. Glavina J, Roman EA, Espada R, de Prat-Gay G, Chemes LB, Sanchez IE. Interplay between sequence, structure and linear motifs in the adenovirus E1A hub protein. Virology. 2018;525:117–131. doi:10.1016/j.virol.2018.08.012

44. Lau L, Gray EE, Brunette RL, Stetson DB. DNA tumour virus oncogenes antagonize the cGAS-STING DNA-sensing pathway. Science. 2015;350(6260):568–571. doi:10.1126/science.aab3291

## REFERENCES

45. Chemes LB, Noval MG, Sanchez IE, de Prat-Gay G. Folding of a cyclin box: linking multitarget binding to marginal stability, oligomerization, and aggregation of the retinoblastoma tumour suppressor AB pocket domain. J Biol Chem. 2013;288(26):18923–18938. doi:10.1074/jbc.M113.467316

46. Uversky VN. What does it mean to be natively unfolded? Eur J Biochem. 2002;269(1):2–12. http://www.ncbi.nlm.nih.gov/pubmed/11784292

47. Hofmann H, Soranno A, Borgia A, Gast K, Nettels D, Schuler B. Polymer scaling laws of unfolded and intrinsically disordered proteins quantified with single-molecule spectroscopy. Proc Natl Acad Sci U S A. 2012;109(40):16155–16160. doi:10.1073/pnas.1207719109

48. Kuzmic P, Moss ML, Kofron JL, Rich DH. Fluorescence displacement method for the determination of receptor-ligand binding constants. Anal Biochem. 1992;205(1):65–69. doi:10.1016/0003-2697(92)90579-v

49. Perozzo R, Folkers G, Scapozza L. Thermodynamics of protein-ligand interactions: history, presence, and future aspects. J Recept Signal Transduct Res. 2004;24(1-2):1–52. http://www.ncbi.nlm.nih.gov/pubmed/15344878

50. Muhandiram DR, Kay LE. Gradient-Enhanced Triple-Resonance Three-Dimensional NMR Experiments with Improved Sensitivity. J Magn Reson Ser B. 1994;103(3):203–216. doi:https://doi.org/10.1006/jmrb.1994.1032

51. Wittekind M, Mueller L. HNCACB, a High-Sensitivity 3D NMR Experiment to Correlate Amide-Proton and Nitrogen Resonances with the Alpha- and Beta-Carbon Resonances in Proteins. J Magn Reson Ser B. 1993;101(2):201–205. doi:https://doi.org/10.1006/jmrb.1993.1033

52. Johnson R.A. Bar. B. NMRView: a computer program for the visualization and analysis of NMR data. J Biomol NMR. 1994;4:603–614.

53. Tamiola K, Acar B, Mulder FA. Sequence-specific random coil chemical shifts of intrinsically disordered proteins. J Am Chem Soc. 2010;132(51):18000–18003. doi:10.1021/ja105656t

54. Zhou HX. The affinity-enhancing roles of flexible linkers in two-domain DNA-binding proteins. Biochemistry. 2001;40(50):15069–15073. http://www.ncbi.nlm.nih.gov/pubmed/11735389

55. Lee JO, Russo AA, Pavletich NP. Structure of the retinoblastoma tumour-suppressor pocket domain bound to a peptide from HPV E7. Nature. 1998;391(6670):859–865. doi:10.1038/36038

56. Estaña N.; Delaforge, E.; Vaisset, M.; Cortés, J.; Bernadó, P. A. S. Realistic ensemble models of intrinsically disordered proteins using a structure-encoding coil database.

57. Franke D, Petoukhov M V, Konarev P V, et al. ATSAS 2.8: a comprehensive data analysis suite for small-angle scattering from macromolecular solutions. J Appl Crystallogr. 2017;50(Pt 4):1212–1225. doi:10.1107/S1600576717007786

58. Guinier A. Diffraction of x-rays of very small angles-application to the study of ultramicroscopic phenomenon. Ann Phys. 1939;12:161–237.

59. Svergun Semenyuk,A. V.; Feigin,L.A. DI. Small-angle-scattering-data treatment by the regularization method. Acta Crystallogr Sect A Found Crystallogr. 1988;44:244–250.

60. Balog ER, Burke JR, Hura GL, Rubin SM. Crystal structure of the unliganded retinoblastoma protein pocket domain. Proteins. 2011;79(6):2010–2014. doi:10.1002/prot.23007

61. Schneidman-Duhovny D, Hammel M, Tainer JA, Sali A. Accurate SAXS profile computation and its assessment by contrast variation experiments. Biophys J. 2013;105(4):962–974. doi:10.1016/j.bpj.2013.07.020

62. Schneidman-Duhovny D, Hammel M, Tainer JA, Sali A. FoXS, FoXSDock and MultiFoXS: Single-state and multi-state structural modelling of proteins and their complexes based on SAXS profiles. Nucleic Acids Res. 2016;44(W1):W424–9. doi:10.1093/nar/gkw389

63. Weinkam P, Pons J, Sali A. Structure-based model of allostery predicts coupling between distant sites. Proc Natl Acad Sci U S A. 2012;109(13):4875–4880. doi:10.1073/pnas.1116274109

64. Tria G, Mertens HD, Kachala M, Svergun DI. Advanced ensemble modelling of flexible macromolecules using X-ray solution scattering. IUCrJ. 2015;2(Pt 2):207–217. doi:10.1107/S205225251500202X

65. Svergun Barberato,C.; Koch,M.H.J. D. CRYSOL – a Program to Evaluate X-ray Solution Scattering of Biological Macromolecules from Atomic Coordinates. J Appl Crystallogr. 1995;28:768–773.

66. Garcia De La Torre J, Huertas ML, Carrasco B. Calculation of hydrodynamic properties of globular proteins from their atomic-level structure. Biophys J. 2000;78(2):719–730. doi:10.1016/S0006-3495(00)76630-6

67. Ortega A, Amoros D, Garcia de la Torre J. Prediction of hydrodynamic and other solution properties of rigid proteins from atomic- and residue-level models. Biophys J. 2011;101(4):892–898. doi:10.1016/j.bpj.2011.06.046

68. Vitalis A, Pappu R V. ABSINTH: a new continuum solvation model for simulations of polypeptides in aqueous solutions. J Comput Chem. 2009;30(5):673–699. doi:10.1002/jcc.21005

69. Vitalis A, Pappu R V. Methods for Monte Carlo simulations of biomacromolecules. Annu Rep Comput Chem. 2009;5:49–76. doi:10.1016/S1574-1400(09)00503-9

70. Kozlov AG, Weiland E, Mittal A, et al. Intrinsically disordered C-terminal tails of E. coli single-stranded DNA binding protein regulate cooperative binding to single-stranded DNA. J Mol Biol. 2015;427(4):763–774. doi:10.1016/j.jmb.2014.12.020

71. Metskas LA, Rhoades E. Conformation and Dynamics of the Troponin I C-Terminal Domain: Combining Single-Molecule and Computational Approaches for a Disordered Protein Region. J Am Chem Soc. 2015;137(37):11962–11969. doi:10.1021/jacs.5b04471

72. McGibbon RT, Beauchamp KA, Harrigan MP, et al. MDTraj: A Modern Open Library for the Analysis of Molecular Dynamics Trajectories. Biophys J. 2015;109(8):1528–1532. doi:10.1016/j.bpj.2015.08.015

73. Holehouse AS, Das RK, Ahad JN, Richardson MO, Pappu R V. CIDER: Resources to Analyse Sequence-Ensemble Relationships of Intrinsically Disordered Proteins. Biophys J. 2017;112(1):16–21. doi:10.1016/j.bpj.2016.11.3200

74. Schymkowitz J, Borg J, Stricher F, Nys R, Rousseau F, Serrano L. The FoldX web server: an online force field. Nucleic Acids Res. 2005;33(Web Server issue):W382–8. doi:10.1093/nar/gki387

75. London N, Raveh B, Cohen E, Fathi G, Schueler-Furman O. Rosetta FlexPepDock web server--high resolution modelling of peptide-protein interactions. Nucleic Acids Res. 2011;39(Web Server issue):W249–53. doi:10.1093/nar/gkr431

76. Good P. Permutation, Parametric, and Bootstrap Tests of Hypotheses. 3rd ed. Springer-Verlag New York; 2005. doi:10.1007/b138696

77. Benjamini Y, Hochberg Y. Controlling the False Discovery Rate: a Practical and Powerful Approach to Multiple Testing. J R Stat Soc Ser B. 1995;57(1):289–300. doi:10.2307/2346101

