## EXTENDED AND SUPPLEMENTAL DATA for "Conformational buffering underlies functional selection in intrinsically disordered protein regions": 02 EXTENDED DATA_FOUTEL.pdf

***Extended Data Table 1**: Thermodynamic Parameters of the interaction between Rb and cellular E2F2 and E1A peptides and proteins.*

| | ITC [2] | | | | | Fluorescence Spectroscopy | | | |
|---|---|---|---|---|---|---|---|---|---|
| | | | | | | Direct titration [3] | | Competition [4] | |
| **Titrant [1]** | **n** | ***ΔH*** (cal/mol) | ***-TΔS*** (cal/mol) | ***ΔG*** (cal/mol) | **$K_D$** (nM) | ***ΔG*** (cal/mol) | **$K_D$** (nM) | **$K_D$** (nM) | **IC50** (nM) |
| **E2F2** | 0.94 ± 0.01 | -8646 ± 174 | -2170 ± 302 | -10817 ± 247 | 8.7 ± 3.6 | -12057 ± 16 | 1.03 ± 0.03 | 4.7 ± 0.3 | 30 |
| **E1A$_{E2F}$** | 0.73 ± 0.06 | 1100 ± 144 | -9102 ± 469 | -8003 ± 446 | 1080 ± 828 | -9286 ± 10 | 119.1 ± 2.1 | 299 ± 45 | 2000 |
| **E1A$_{LxCxE}$** | 1.03 ± 0.01 | -9522 ± 79 | 573 ± 92 | -8949 ± 47 | 213 ± 17 | -9362 ± 16 | 104.6 ± 2.9 | - | - |
| **E1A$_{LxCxE\text{-AC}}$** | 1.02 ± 0.01 | -10127 ±122 | 511 ± 161 | -9615 ± 104 | 68.5 ± 12.3 | -9577 ± 18 | 73.4 ± 2.2 | *nd* | *nd* |
| **E1A$_{LxCxE\text{-ACP}}$** | 1.19 ± 0.01 | -7900 ± 107 | -2021 ± 168 | -9921 ± 129 | 40.6 ± 9.1 | -10339 ± 16 | 19.6 ± 0.5 | *nd* | *nd* |
| **E1A$_{WT}$** | 1.19 ± 0.01 | -5284 ± 40 | -8960 ± 105 [a] | *nd* | *nd* | -14244 ± 97 | 0.024 ± 0.004 | 0.065 ± 0.009 | 1.5 |
| **E1A$_{ΔE}$** | 0.92 ± 0.01 | -10784 ± 95 | 452 ± 140 | -10332 ± 103 | 19.9 ± 3.8 | -9828 ± 21 | 47.1 ± 1.7 | *nd* | *nd* |
| **E1A$_{ΔL}$** | 0.92 ± 0.05 | 2447 ± 291 | -10553 ± 477 | -8106 ± 378 | 904 ± 578 | -8966 ± 9 | 206.7 ± 3.1 | 248 ± 37 | 1500 |

Most measurements were performed in triplicate except for the ITC titrations for Rb-E1A$_{WT}$ which was performed in duplicate and for Rb-E1A$_{E2F}$ which was performed once. For competition experiments, titrations were performed at least twice but values are from a single experimental set.

[1] All ITC and Fluorescence Spectroscopy measurements were performed at 20.0 °C.

[2] For ITC experiments, n values are the average of all independent experimental determinations and the standard deviation is obtained as the propagated mean standard error. *ΔH* and $K_D$ values are the average of all independent experimental determinations. The standard deviations for *ΔH* and $K_D$ are obtained as propagated mean standard errors. *ΔG* is calculated as $ΔG = RT{*}LnK_D$ and *-TΔS* as $-TΔS = ΔG - ΔH$. The standard deviations for *ΔG* and *-TΔS* are obtained as the propagated mean standard errors.

[3] For Fluorescence Spectroscopy direct titration experiments, the $K_D$ value was obtained from a global fitting of three independent experimental determinations performed at different protein concentrations (See materials and methods). The error for $K_D$ is the standard error of the fitted parameter. *ΔG* is calculated as $ΔG = RT{*}LnK_D$ and its standard deviation is obtained by error propagation.

[4] For Fluorescence Spectroscopy competition experiments, the error for $K_D$ is the standard error of the fitted parameter.

[a] Calculated using *ΔG* from Fluorescence data

Values for thermodynamic parameters are: R = 1.985 cal/K*mol. T = 283.15 K (10 ºC) or T = 293.15 K (20 ºC).

- No value could be fitted to the displacement curve as it had no change in fluorescence over the concentration range tested.

*nd:* not determined

***Extended Data Table 2**: Dissociation constants for the interaction between Rb and cellular E2F2 and E1A peptides and proteins*

| **Probe** | (nM) | $K_D$ (nM) | $K_D$ (nM) | **r**(*0*) | **r**(*F*) | **F**(*0*) | **F**(*F*) |
|---|---|---|---|---|---|---|---|
| | | Raw data | Normalized data | | | | |
| **E2F2** | 1 | 1.18 ± 0.16 | 1.15 ± 0.05 | 0.062 | 0.163 | | |
| | 5 | 0.91 ± 0.04 | 0.91 ± 0.02 | 0.049 | 0.209 | | |
| | 10 | 0.71 ± 0.05 | 0.74 ± 0.03 | 0.043 | 0.219 | | |
| | 20 | 1.39 ± 0.16 | 1.55 ± 0.05 | 0.043 | 0.215 | | |
| | 30 | 1.06 ± 0.08 | 1.14 ± 0.05 | 0.043 | 0.227 | | |
| | $K_D$ global fit | | 1.03 ± 0.03 | | | | |
| **E1A$_{E2F}$** | 100 | 109.3 ± 9.7 | 97.6 ± 3.6 | 0.046 | 0.198 | | |
| | 200 | 100.2 ± 22.4 | 97.4 ± 5.6 | 0.049 | 0.206 | | |
| | 300 | 101.3 ± 11.4 | 93.2 ± 4.2 | 0.049 | 0.206 | | |
| | 500 | 105.5 ± 10.2 | 93.4 ± 4.4 | 0.050 | 0.211 | | |
| | 1000 | 127.9 ± 12.2 | 75.1 ± 7.9 | 0.045 | 0.203 | | |
| | 2000 | 109.5 ± 0.6 | 81.6 ± 4.9 | 0.046 | 0.206 | | |
| | $K_D$ global fit | | 119.1 ± 2.1 | | | | |
| **E1A$_{LxCxE}$** | 100 | 113.4 ± 14.1 | 117.2 ± 6.5 | 0.028 | 0.076 | | |
| | 200 | 93.1 ± 11.3 | 93.5 ± 2.9 | 0.036 | 0.091 | | |
| | 300 | 93.7 ± 7.9 | 93.0 ± 2.6 | 0.033 | 0.087 | | |
| | 500 | 97.9 ± 15.6 | 98.7 ± 3.7 | 0.032 | 0.088 | | |
| | 1000 | 117.1 ± 28.3 | 111.9 ± 7.4 | 0.032 | 0.086 | | |
| | $K_D$ global fit | | 104.6 ± 2.9 | | | | |
| **E1A$_{LxCxE\text{-}AC}$** | 130 | 80.7 ± 12.8 | 80.5 ± 6.9 | 0.04 | 0.100 | | |
| | 175 | 73.7 ± 5.2 | 73.7 ± 5.2 | 0.04 | 0.105 | | |
| | 700 | 85.6 ± 7.4 | 82.8 ± 5.8 | 0.039 | 0.106 | | |
| | $K_D$ global fit | | 73.4 ± 2.2 | | | | |
| **E1A$_{LxCxE\text{-}ACP}$** | 30 | 20.1 ± 2.6 | 20.1 ± 2.6 | 0.05 | 0.099 | | |
| | 50 | 20.2 ± 2.4 | 17.5 ± 1.3 | 0.04 | 0.093 | | |
| | 100 | 19.5 ± 2.2 | 18.2 ± 2.2 | 0.039 | 0.092 | | |
| | $K_D$ global fit | | 19.6 ± 0.5 | | | | |
| **E1A$_{WT}$** | 0.5 | 0.025 ± 0.004 | 0.025 ± 0.003 | 0.075 | 0.191 | | |
| | 0.5 | 0.027 ± 0.006 | 0.027 ± 0.003 | 0.059 | 0.127 | | |
| | 0.5 | 0.026 ± 0.009 | 0.025 ± 0.008 | | | 111.6 | 94.3 |
| | 1 | 0.037 ± 0.009 | 0.034 ± 0.005 | 0.061 | 0.153 | | |
| | 1 | 0.049 ± 0.019 | 0.050 ± 0.012 | | | 302.8 | 259.8 |
| | 2 | 0.046 ± 0.007 | 0.047 ± 0.005 | 0.065 | 0.197 | | |
| | 2 | 0.062 ± 0.008 | 0.062 ± 0.005 | 0.064 | 0.2 | | |
| | 2 | 0.065 ± 0.015 | 0.062 ± 0.009 | 0.062 | 0.175 | | |
| | 2 | 0.065 ± 0.022 | 0.092 ± 0.025 | | | 451.3 | 374.3 |
| | 2 | 0.074 ± 0.036 | 0.090 ± 0.034 | | | 412.4 | 336.6 |
| | 2 | 0.075 ± 0.010 | 0.074 ± 0.006 | | | 613.5 | 503.4 |
| | $K_D$ global fit | | 0.024 ± 0.004 | | | | |
| **E1A$_{\Delta E}$** | 50 | 42.6 ± 3.9 | 42.5 ± 3.6 | 0.05 | 0.105 | | |
| | 200 | 53.9 ± 3.1 | 53.9 ± 3.1 | 0.05 | 0.106 | | |
| | 800 | 60.2 ± 10.1 | 54.5 ± 10.3 | 0.048 | 0.105 | | |
| | $K_D$ global fit | | 47.1± 1.7 | | | | |
| **E1A$_{\Delta L}$** | 200 | 197.7 ± 4.5 | 197.7 ± 4.5 | 0.058 | 0.220 | | |
| | 400 | 207.5 ± 7.4 | 207.5 ± 7.4 | 0.056 | 0.232 | | |
| | 800 | 225.2 ± 20.6 | 217.9 ± 10.9 | 0.023 | 0.219 | | |
| | $K_D$ global fit | | 206.7 ± 3.1 | | | | |

Fluorescence spectroscopy direct titrations were performed at fixed concentrations of FITC-labeled protein/peptide titrated with increasing amounts of Rb until saturation. For each complex this experiment was performed at different concentrations of FITC-probes. The $K_D$ parameter was obtained from fitting the titration curves to a bimolecular association model from individual titrations using raw and normalized titration curves or by global fitting of the normalized data. An excellent agreement between individual and global fits was obtained. Initial anisotropy signals *r(0)*, showed that intrinsic anisotropy of the FITC-probes are constant and independent of the concentrations tested. Final anisotropy signals *r(F)* reached stable values for the complex with Rb. Initial and final fluorescence signals (*F(0) y F(F)*) were measured only for E1A$_{WT}$ and even though these values represent a more variable range, they proved to be equally useful for fitting $K_D$ values.

***Extended Data Table 3**: Analysis of quaternary structure and hydrodynamic behavior of unbound Rb and E1A proteins and [Rb:E1A] complexes.*

| | $MW_{THEO}$ [1] (kDa) | $MW_{SLS}$ [2] (kDa) | $MW_{SLS}$ [3] $/MW_{THEO}$ | $MW_{app\ SEC}$ [4] $/MW_{THEO}$ | $R_h$ (nm) [5] EXPERIMENTAL | $R_h$ (nm) [6] ENSEMBLE 25ºC | $R_h$ (nm) [7] ENSEMBLE 20ºC | $R_g$ (nm) [8] SAXS | $R_g/R_h$ [9] | v [10] |
|---|---|---|---|---|---|---|---|---|---|---|
| **E1A$_{WT}$** | 12.5 | 13.93 | 1.11 | 4.35<br>4.77 ± 0.01[a] | 3.07 ± 0.12<br>3.42 ± 0.01[b] | - | - | 4.28 ± 0.02<br>4.64 ± 0.03<br>4.51 ± 0.02 | 1.39<br>1.25[d] | 0.64[e]<br>0.53[f]<br>0.55[g] |
| **E1A$_{\Delta L}$** | 12.2 | 11.58 | 0.95 | 4.40 | 3.03 ± 0.12 | - | - | - | - | - |
| **E1A$_{\Delta E}$** | 12.1 | 12.96 | 1.07 | 4.36 | 3.07 ± 0.12 | - | - | - | - | - |
| **Rb** | 42.1 | 38.82 | 0.92 | 1.09 | 2.94 ± 0.12 | - | - | 2.42 ± 0.02<br>2.50 ± 0.01<br>2.61 ± 0.01 | 0.82 | 0.38[f] |
| **[Rb:E1A$_{WT}$]** | 54.6 | 56.3 | 1.03 | 1.12 | 3.20 ± 0.12 | 3.27<br>3.36[c] | 3.71<br>3.79[c] | 2.98 ± 0.04<br>2.97 ± 0.02<br>3.33 ± 0.02 | 0.93 | - |
| **[Rb:E1A$_{\Delta E}$]** | 54.2 | 53.9 | 0.99 | 1.20 | 3.34 ± 0.13 | 3.64 | 4.12 | - | - | - |
| **[Rb:E1A$_{\Delta L}$]** | 54.3 | 56.2 | 1.03 | 1.27 | 3.27 ± 0.12 | 3.60 | 4.07 | - | - | - |

[1] $MW_{THEO}$ is the theoretical monomeric molecular weight of each free protein and for the complexes it results from the sum of their molecular weights considering a 1:1 complex.
[2] $MW_{SLS}$ was experimentally determined by SEC-SLS at RT.
[3] $MW_{SLS}/MW_{THEO}$ ratios indicate the oligomerization state of unbound proteins and stoichiometry for the protein complexes. Value 1 indicates a monomer state for unbound proteins and a [1:1] stoichiometry for complexes.
[4] $MW_{appSEC}/MW_{THEO}$ ratios indicate the extended or compact hydrodynamic behavior of the unbound proteins and complexes. $MW_{appSEC}$ corresponds to the molecular weight estimated from the calibration curve in SEC experiments (**See Methods**). Values above 1 indicate an extended behavior.
[5] $R_{h\ EXPERIMENTAL}$ was calculated from SEC data as: log $R_h$ = -0.254 + 0.369 log $MW_{appSEC}$ [[1]] and its standard deviation was obtained by error propagation.
[6,7] $R_{h\ ENSEMBLE}$ was calculated for generated conformations at 25 ºC and 20 ºC respectively using the program HYDROPRO as explained in (**See Methods**) or from the sub-ensembles selected by EOM.
[8] $R_{g\ SAXS}$ was calculated from SAXS experiments. SAXS measurements were performed at three different concentrations for E1A$_{WT}$, Rb and the [Rb:E1A$_{WT}$] complex (**See Methods**) and for the sake of completeness $R_g$ derived from all concentrations tested are informed. An increase in $R_g$ at 2.7 mg/ml of [Rb:E1A$_{WT}$] complex indicated inter-particle interaction and the Guinier region for this concentration was not used for further analysis.
[9] Ratio of SAXS-derived $R_g$ to experimental $R_h$ ($R_g/R_h$) for free and bound proteins. The $R_g$ values used here correspond to those calculated from the Guinier region in the final merged SAXS profiles of E1A$_{WT}$, Rb and [Rb:E1A$_{WT}$] complex, which contain information from the lowest concentration data. Ratio values of 0.75, 1.0 and 1.50 are expected for compact globules, Flory random chains (FRC) and for excluded volume (EV) chains respectively. For Flory random chains, modest deviations around 1.0 are probably the result of stronger intrachain repulsions or attractions [[2]]. Unless noted otherwise the $R_h$ values used here are derived from SEC experiments.
[10] The exponent $v$ was calculated by applying the scaling law $R_x=R_0N^v$ where $R_x$ might be $R_h$ or $R_g$. and $R_0$ depends on the dimensions of the chain and was obtained as described in e,f and g. The reference values for v depend on the dimensions of the chain and are 0.6 for an excluded volume chain, 0.33 for a compact globule and 0.5 for a chain in the Θ-regime [[3]].
[a] Found as $MW_{DLS}/MW_{THEO}$ where $MW_{DLS}$ was obtained from three independent DLS measurements (**See Methods**).
[b] Obtained from three independent DLS measurements.
[c] $R_h$ values corresponding to the sub-ensembles selected by EOM that best fit SAXS data (**Figure 3**).
[d] $R_g/R_h$ for E1A$_{WT}$ using $R_h$ derived from DLS experiments.
[e] v was fitted using $R_g$ obtained from SAXS measurements using the formula $R_g=R_0N^v$ with $R_0$ = 2.1 nm and N= 114 for E1A$_{WT}$. The $R_0$ value is from [[3]].
[f] v was fitted using $R_h$ obtained from SEC-SLS measurements using the formula $R_h=R_0N^v$ with $R_0$ = 2.49 nm and N= 114 for E1A$_{WT}$ and $R_0$ = 4.92 nm and N= 360 for Rb. The $R_0$ values are from [[4]].
[g] v was fitted using $R_h$ obtained from using DLS measurements using the formula $R_h=R_0N^v$ with $R_0$ = 2.49 nm and N= 114 for E1A$_{WT}$. The $R_0$ values are from [[4]].

***Extended Data Table 4****: Structural parameters from calorimetric data for the interactions between Rb and* E1A$_{\Delta L}$ and E1A$_{E2F}$*.*

| Titrant | °C | n [1] | $\Delta H$ (cal/mol) | $-T\Delta S$ (cal/mol) | $\Delta G$ (cal/mol) | $K_D$ (nM) | $\Delta Cp$ (cal/mol K) | $\Delta ASA_T$ [2] ($Å^2$) | $X_{res}$ [3] |
|---|---|---|---|---|---|---|---|---|---|
| **E1A$_{\Delta L}$** | 10 | 0.91 ± 0.03 | 8675 ± 599 | -16196 ± 622 | -7521 ± 166 | 1590 ± 487 | | | |
| **E1A$_{\Delta L}$** | 20 | 0.92 ± 0.05 | 2447 ± 291 | -10553 ± 477 | -8106 ± 378 | 904 ± 578 | -606.3 ± 9.6 | -1854 ± 55 | 28 ± 1 |
| **E1A$_{\Delta L}$** | 30 | 0.94 ± 0.03 | -3450 ± 225 | -4920 ± 310 | -8379 ± 213 | 922 ± 326 | | | |
| **E1A$_{E2F}$** | 10 | 0.86 ± 0.02 | 7205 ± 338 | -14739 ± 364 | -7534 ± 134 | 1535 ± 391 | | | |
| **E1A$_{E2F}$** | 15 | 0.86 ± 0.01 | 4130 ± 122 | -11935 ± 156 | -7806 ± 98 | 1200 ± 205 | -629.4 ± 7.3 | -2016 ± 47 | 31± 1 |
| **E1A$_{E2F}$** | 20 | 0.73 ± 0.06 | 1100 ± 144 | -9102 ± 469 | -8003 ± 446 | 1080 ± 828 | | | |
| **E1A$_{E2F}$** | 30 | 0.85 ± 0.02 | -5380 ± 246 | -2960 ± 301 | -8341 ± 173 | 969 ± 279 | | | |

The $\Delta H$ and $K_D$ values are the average of all independent experimental determinations. The errors for $\Delta H$ and $K_D$ are obtained as propagated mean standard errors. $\Delta G$ is calculated as $\Delta G = RT*LnK_D$ and $-T\Delta S$ as $-T\Delta S = \Delta G - \Delta H$. The errors for $\Delta G$ and $-T\Delta S$ are obtained as the propagated mean standard errors.
Values for thermodynamic parameters are: R = 1.985 cal/K*mol. T = 283.15 K (10 ºC), T = 288.15 K (15 ºC), T= 293.15 K (20 ºC) and T = 303.15 K (30 ºC) .
$\Delta C_p$, $\Delta ASA_T$ and $X_{res}$ values were calculated as described in [[5]].
[1] n values are the average of all independent experimental determinations and the standard deviation is obtained as the propagated mean standard error.
[2] According to structural data (PDB 2R7G [[6]]), the formation of the [Rb:E1A$_{E2F}$] complex causes a total burial of accessible surface area of $\Delta ASA_{total}$ = 1623 $Å^2$, in close agreement with our estimates from ITC.
[3] $X_{res}$ is the number of residues involved in an interaction. Based on a criterion of 4$Å$ separation between residues (upper limit for non-covalent interactions), the structural analysis yielded 32 residues (23 from Rb and 9 from E1A$_{E2F\,(37\text{-}49)}$) involved in the interaction, in close agreement with our estimates from ITC.

***Extended Data Table 5***: *Analysis of the allosteric effect between E1A binding sites.*

| Cell | Titrant [1] | n [2] | $\Delta H$ (cal/mol) | $-T\Delta S$ (cal/mol) | $\Delta G$ (cal/mol) | $K_D$ (nM) | $\Delta\Delta G$ [3] (cal/mol) |
|---|---|---|---|---|---|---|---|
| **Rb** | **E1A$_{E2F}$** | 0.86 ± 0.02 | 7205 ± 338 | -14739 ± 364 | -7534 ± 134 | 1535 ± 392 | |
| **Rb + E1A$_{LxCxE}$** | **E1A$_{E2F}$** | 0.89 ± 0.01 | 6875 ± 214 | -14510 ± 236 | -7634 ± 99 | 1327 ± 159 | -100 ± 166 |
| **Rb** | **E1A$_{\Delta L}$** | 0.91 ± 0.03 | 8675 ± 599 | -16196 ± 622 | -7521 ± 166 | 1590 ± 487 | |
| **Rb + E1A$_{LxCxE}$** | **E1A$_{\Delta L}$** | 0.86 ± 0.02 | 6795 ± 320 | -14572 ± 351 | -7777 ± 144 | 994 ± 252 | -256 ± 220 |
| **Rb** | **E1A$_{LxCxE}$** | 1.03 ± 0.01 | -9522 ± 79 | 573 ± 92 | -8949 ± 47 | 213 ± 17 | |
| **Rb + E1A$_{E2F}$** | **E1A$_{LxCxE}$** | 1.19 ± 0.01 | -8301 ± 89 | -565 ± 110 | -8867 ± 37 | 245 ± 27 | 82 ± 60 |
| **Rb + E1A$_{\Delta L}$** | **E1A$_{LxCxE}$** | 1.13 ± 0.01 | -8873 ± 64 | 78 ± 76 | -8795 ± 41 | 277 ± 20 | 154 ± 62 |

All measurements were performed in triplicate except for the Rb-E1A$_{E2F}$, (Rb+E1A$_{LxCxE}$)-E1A$_{E2F}$ and (Rb+E1A$_{LxCxE}$)-E1A$_{\Delta L}$ titrations, which were performed in duplicate.

[1] Titrations with E1A$_{E2F}$ and E1A$_{\Delta L}$ as titrants were performed at 10.0 °C and titrations with E1A$_{LxCxE}$ as titrant, were performed at 20.0 °C.

[2] n values are the average of all independent experimental determinations and the standard deviation is obtained as the propagated mean standard error.

[3] $\Delta\Delta G$ was calculated as $\Delta\Delta G = \Delta G$ pre-saturated - $\Delta G$ non-saturated with the complementary motif (or region) to the titrant.

The $\Delta H$ and $K_D$ values reported are the average of all independent experimental determinations. The standard deviations for $\Delta H$ and $K_D$ are obtained as propagated mean standard errors. $\Delta G$ is calculated as $\Delta G = RT*LnK_D$ and $-T\Delta S$ as $-T\Delta S = \Delta G - \Delta H$. The standard deviations for $\Delta G$ and $-T\Delta S$ are obtained as the propagated mean standard errors.

Values for thermodynamic parameters are: R = 1.985 cal/K*mol. T = 283.15 K (10 ºC) or T = 293.15 K (20 ºC).

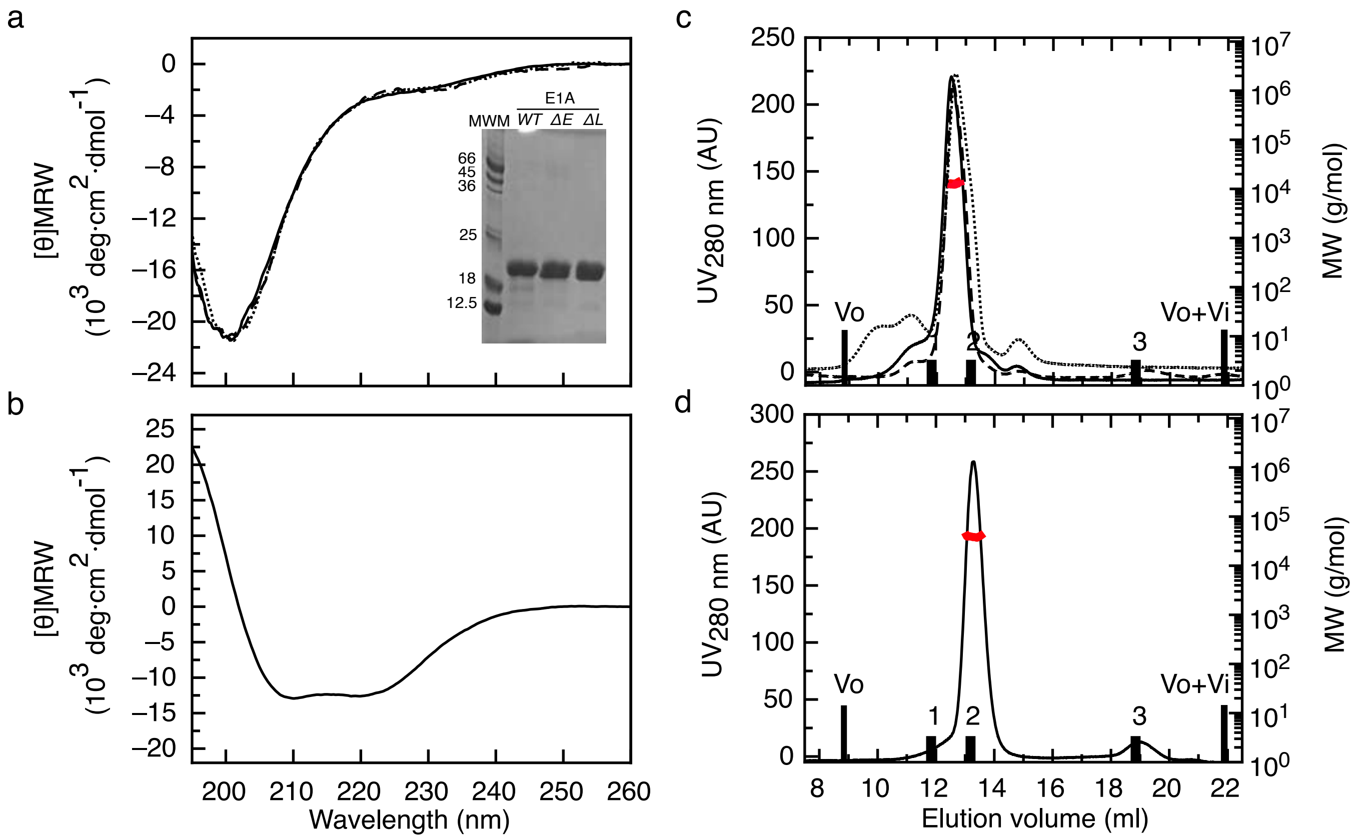

a
b
c
d
[θ]MRW
(10^3 deg·cm^2·dmol^-1)
E1A
MWM
WT
ΔE
ΔL
66
45
36
25
18
12.5
Wavelength (nm)
UV280 nm (AU)
MW (g/mol)
Elution volume (ml)
Vo
Vo+Vi
1
2
3

**EXTENDED DATA FIGURE 1: Biophysical characterization of recombinant Rb and E1A proteins.**
**a)** Far UV-CD spectra of E1A$_{WT}$ (solid line), E1A$_{\Delta E}$ (dotted line), E1A$_{\Delta L}$ (dashed line). Inset: 15% SDS-PAGE gel of purified recombinant E1A proteins (purity > 90%). **b)** Far UV-CD spectrum of the RbAB domain. **c)** SEC-SLS experiments of E1A$_{WT}$ (solid line), E1A$_{\Delta E}$ (dotted line) and E1A$_{\Delta L}$ (dashed line). **d)** SEC-SLS experiment of the RbAB domain. For b) and c), black bars correspond to the exclusion volume of globular protein markers: BSA 66 kDa (1), MBP 45 kDa (2) and Lysozyme 14.3 kDa (3). Black line: SEC profile, red line: measurement of the molecular weight.

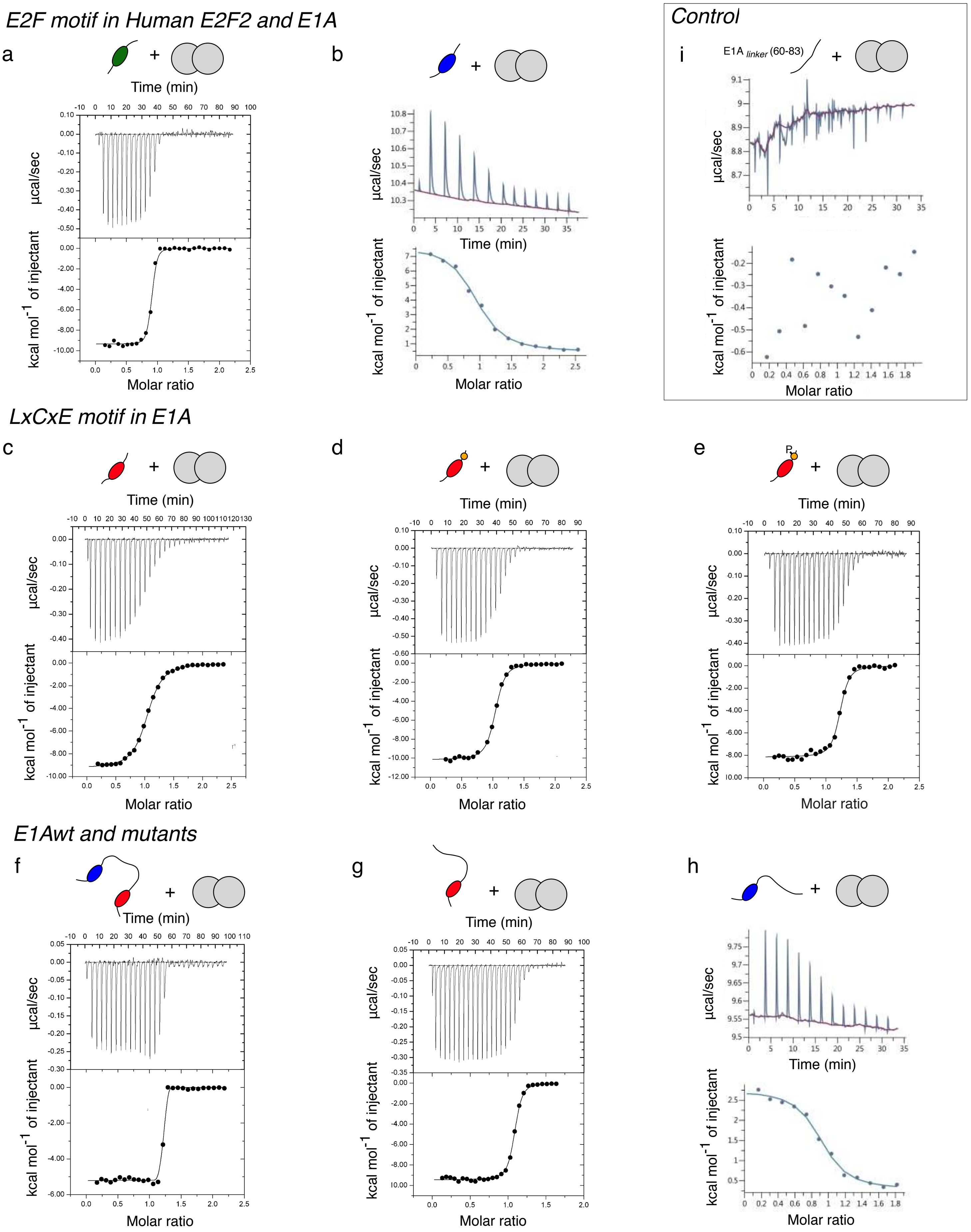

E2F motif in Human E2F2 and E1A
a
b
Control
i
E1A linker (60-83)
LxCxE motif in E1A
c
d
e
P
E1Awt and mutants
f
g
h
Time (min)
µcal/sec
kcal mol-1 of injectant
Molar ratio

**EXTENDED DATA FIGURE 2: Isothermal Titration Calorimetry experiments.**

Representative ITC binding curves for each Rb: peptide/protein tested in this work. Measurements were performed loading the cell with Rb solution and the syringe with the different peptides or proteins as titrants. Panels show heat exchanged as a function of time (upper panel), and the enthalpy per mole of injectant plotted as a function of [peptide/protein]/[Rb] molar ratio (lower panel, black circles) and the corresponding fit using a single site binding model (lower panel, black lines). Binding traces here represented correspond to: **a)** Rb (5 µM) and Human E2F2 (50 µM); **b)** Rb (30 µM) and $E1A_{E2F}$ (300 µM); **c)** Rb (15 µM) and $E1A_{LxCxE}$ (150 µM); **d)** Rb (15 µM) and $E1A_{LxCxE\text{-}AC}$ (150 µM); **e)** Rb (15 µM) and $E1A_{LxCxE\text{-}ACP}$ (150 µM); **f)** Rb (15 µM) and $E1A_{WT}$ (150 µM); **g)** Rb (15 µM) and $E1A_{\Delta E}$ (150 µM); **h)** Rb (30 µM) and $E1A_{\Delta L}$ (300 µM). Thermodynamic parameters derived from the fitting are shown in **Extended data table 1.** Exothermic binding to Rb was observed for the Cellular E2F2 peptide and E1A peptides and protein fragments harboring the LxCxE motif, while $E1A_{E2F}$ and $E1A_{\Delta L}$ harboring only the E1A E2F motif clearly showed an endothermic behavior (and required higher concentrations to measure enough heat exchanged). For $E1A_{WT}$ harboring both motifs, there was a compensatory effect with predominant exothermic behavior in the interaction with Rb and a result of their high affinity, $\Delta G$ and $K_D$ could not be determined with this technique (**Extended data table 1**). **i)** To discard an interaction between the E1A linker and Rb, a series of ITC titrations were performed by using a peptide corresponding to TAZ-2 minimal binding region (63-80) in the E1A linker., which showed intensity decreases in the NMR experiments (**Figure 2**). The titration was performed at 30 µM Rb and 300 µM E1A linker peptide at 20 ºC. As it shown, there was no heat exchange upon titration of Rb with E1A linker peptide ruling out a direct interaction. A schematic representation of each interacting pair is shown above the ITC traces: Rb (grey double circle) and each peptide/protein, where binding motifs are represented as follows: Human-E2F2 (green oval), E2F motif (blue oval), LxCxE motif (red oval), LxCxE acidic stretch (orange circle), phosphorylation (letter P). The linker is represented by a black line.

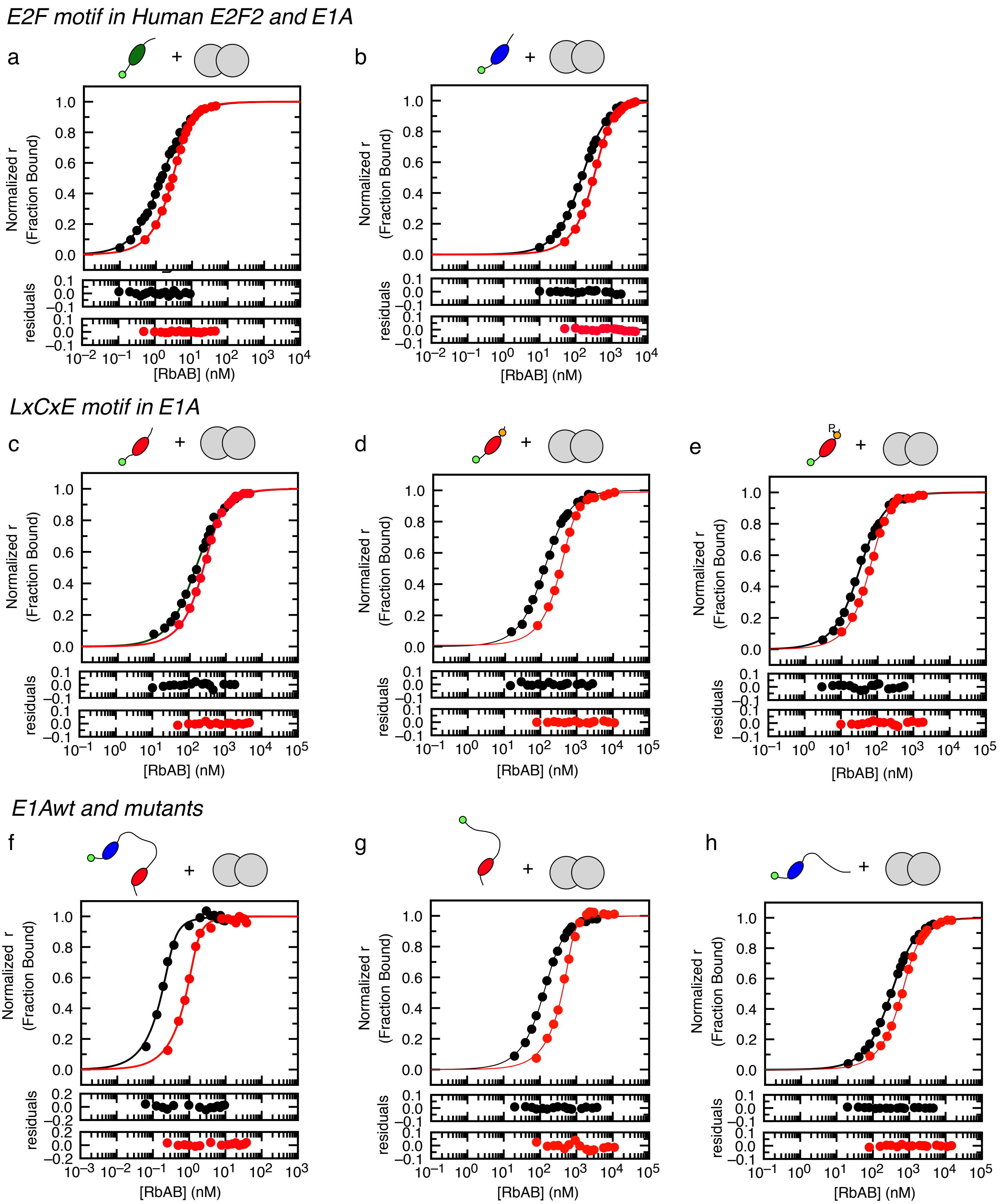

E2F motif in Human E2F2 and E1A
a
b
LxCxE motif in E1A
c
d
e
P
E1Awt and mutants
f
g
h
+
Normalized r (Fraction Bound)
residuals
[RbAB] (nM)

**EXTENDED DATA FIGURE 3: Fluorescence Spectroscopy titration experiments of E1A-Rb and E2F-Rb interactions.**

Representative titration binding curves at equilibrium for each FITC-labeled peptide/protein-Rb interaction tested in this work. Normalized anisotropy signals (circles) are shown, along with the global fit to a 1:1 binding model (lines) that yielded the $K_D$ value. The residuals for the fit are shown in the lower panels. Binding traces here represented correspond to two probe (FITC-labeled peptide/protein) concentrations: **a)** Cellular E2F2: 1 nM (black) and 5 nM (red); **b)** E1A$_{E2F}$: 100 nM (black) and 500 nM (red); **c)** E1A$_{LxCxE}$: 100 nM (black) and 500 nM (red); **d)** E1A $_{LxCxE\text{-}AC}$: 130 nM (black) and 700 nM (red); **e)** E1A $_{LxCxE\text{-}ACP}$: 30 nM (black) and 100 nM (red); **f)** E1A$_{WT}$: 0.5 nM (black) and 2 nM (red); **g)** E1A$_{\Delta E}$: 200 nM (black) and 800 nM (red); **h)** E1A$_{\Delta L}$: 200 nM (black) and 800 nM (red). The $K_D$ values obtained by global fitting to a 1:1 model (**Extended Data Table 1**) were in excellent agreement with those obtained when fitting individual binding curves using non-normalized anisotropy or fluorescence data (**Extended Data Table 2**). A schematic representation of each interacting pair is shown above the binding traces: Rb (grey double circle); FITC-moiety at the N-terminus of the sequence (light green circle). Binding motifs are represented as follows: Human-E2F2 (green oval), E2F motif (blue oval), LxCxE motif (red oval), acidic stretch (orange circle), phosphorylation (letter P). The linker is represented by a black line.

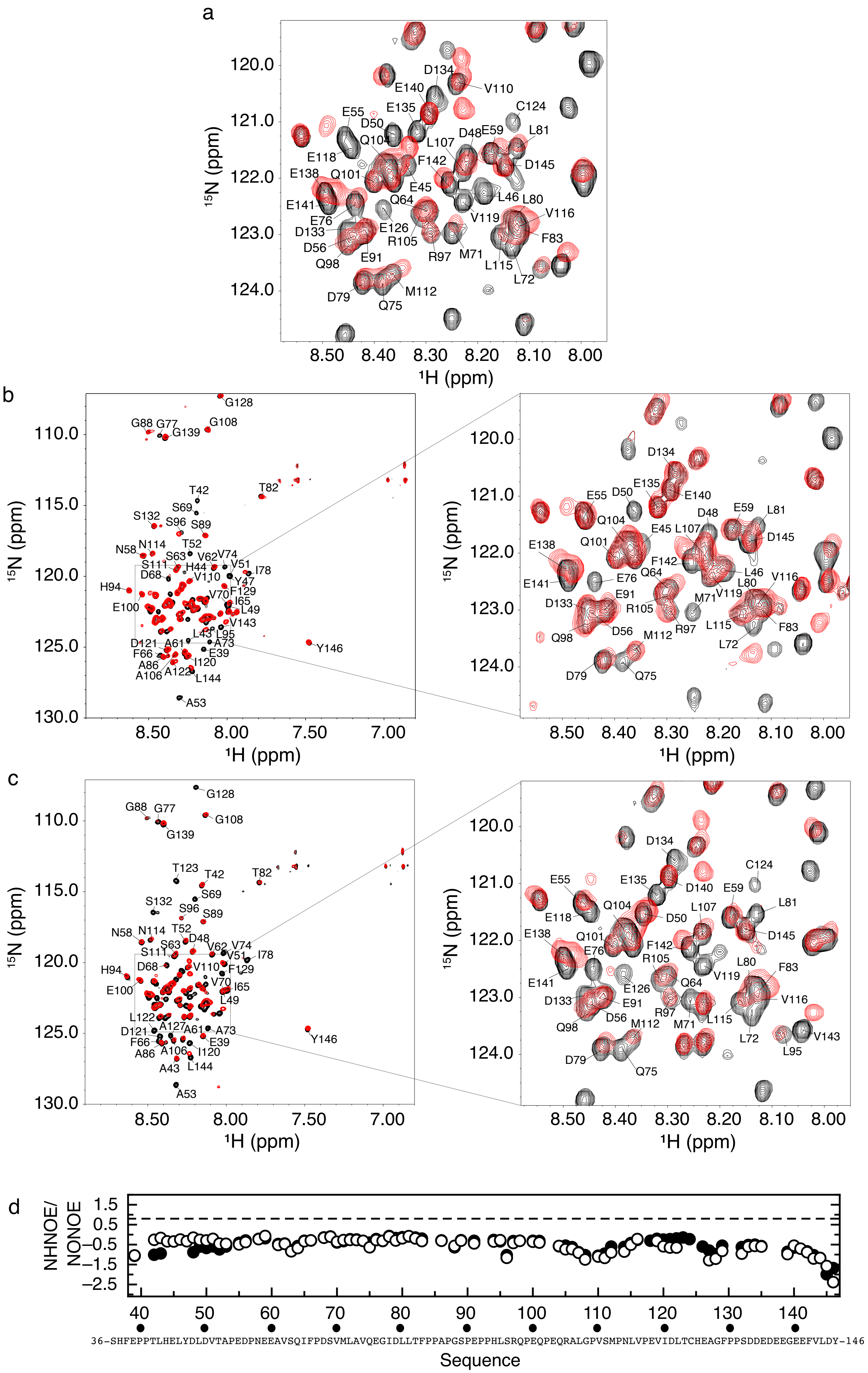
a
120.0
121.0
122.0
123.0
124.0
15N (ppm)
8.50
8.40
8.30
8.20
8.10
8.00
1H (ppm)
D134
E140
V110
E135
C124
E55
D50
Q104
L107
D48
E59
L81
E118
F142
D145
E138
Q101
E45
L46
L80
E141
Q64
V119
V116
E76
D133
E126
R105
F83
D56
R97
M71
Q98
E91
L115
L72
D79
M112
Q75
b
G128
G88
G77
G108
G139
T82
T42
S69
S132
S96
S89
N58
N114
T52
S63
V62
V74
S111
H44
V51
I78
D68
V110
Y47
H94
F129
V70
I65
E100
L49
V143
L43
L95
D121
A61
A73
Y146
F66
E39
A86
I120
A106
A122
L144
A53
110.0
115.0
120.0
125.0
130.0
15N (ppm)
8.50
8.00
7.50
7.00
1H (ppm)
120.0
121.0
122.0
123.0
124.0
D134
E135
E55
D50
E140
D48
E59
L81
Q104
E45
L107
E138
Q101
D145
F142
Q64
E141
E76
L46
L80
V116
E91
M71
V119
D133
R105
F83
Q98
D56
M112
R97
L115
L72
D79
Q75
8.50
8.40
8.30
8.20
8.10
8.00
c
G128
G88
G77
G108
G139
T123
T42
T82
S69
S132
S96
S89
N58
N114
T52
D48
S63
V62
V74
S111
V51
I78
H94
D68
V110
F129
E100
V70
I65
L49
L122
D121
A127
A61
A73
Y146
F66
E39
A86
A106
I120
A43
L144
A53
110.0
115.0
120.0
125.0
130.0
15N (ppm)
8.50
8.00
7.50
7.00
1H (ppm)
120.0
121.0
122.0
123.0
124.0
D134
C124
E55
E135
D140
E59
L81
E118
Q104
D50
L107
E138
Q101
D145
E76
F142
R105
E141
E126
Q64
V119
L80
F83
D133
E91
V116
D56
R97
M112
M71
L115
Q98
L72
V143
L95
D79
Q75
8.50
8.40
8.30
8.20
8.10
8.00
d
NHNOE/
NONOE
1.5
0.5
−0.5
−1.5
−2.5
40
50
60
70
80
90
100
110
120
130
140
36-SHFEPPTLHELYDLDVTAPEDPNEEAVSQIFPDSVMLAVQEGIDLLTFPPAPGSPEPPHLSRQPEQPEQRALGPVSMPNLVPEVIDLTCHEAGFPPSDDEDEEGEEFVLDY-146
Sequence

**EXTENDED DATA FIGURE 4: NMR experiments of [Rb:E1A] complexes.**
**a)** Detail of the central region of $^{1}H$-$^{15}N$ TROSY spectra of free $^{15}N$-labeled E1A (black) and a 1:1 molar ratio complex of $^{15}N$-labeled E1A and unlabeled Rb (red) at 525 μM, with assigned peaks of the free form indicated. The full spectrum of this complex is shown in **Fig. 2a. b)** Left panel: Overlay of the $^{1}H$-$^{15}N$ TROSY spectra of free $^{15}N$-labeled E1A$_{\Delta L}$ (black) and a 1:1 molar ratio complex of $^{15}N$-labeled E1A$_{\Delta L}$ and unlabeled Rb (red) at 315 μM. Right panel: detail of the central region of the spectra . with assigned peaks of the free form indicated **c)** Left panel: Overlay of the $^{1}H$-$^{15}N$ TROSY spectra of free $^{15}N$-labeled E1A$_{\Delta E}$ (black) and a 1:1 molar ratio complex of $^{15}N$-labeled E1A$_{\Delta E}$ and unlabeled Rb (red) at 315 μM. Right panel: detail of the central region of the spectra with assigned peaks of the free form indicated. The low chemical shift dispersions in the $^{1}H$-dimension for E1A$_{\Delta L}$ and E1A$_{\Delta E}$ denote their disordered nature, alike that seen in E1A. The lack of increase in peak dispersion upon binding with Rb indicates that E1A$_{\Delta L}$ and E1A$_{\Delta E}$ mutants remain largely disordered in the [Rb:E1A$_{\Delta L}$] and [Rb:E1A$_{\Delta E}$] complexes. **d)** Plot of NHNOE/NONOE ratio as a function of E1A residue number. This method is useful for analyzing the backbone dynamics at pico to nanosecond timescale. Positive values up to +0.8 (dashed line) are typical for rigid polypeptide segments suggesting folded conformations, although this must be interpreted on the light of other parameters such as secondary chemical shifts. Negative NOE values up to -3.5 are indicative of high flexibility which is typical of IDPs. As it can be observed, NOE ratios were below zero across the entire E1A sequence length. N- and C-terminal regions showed values lower than the average caused by their higher flexibility. E1A$_{\Delta L}$ (black empty circles), E1A$_{\Delta E}$ (black circles).

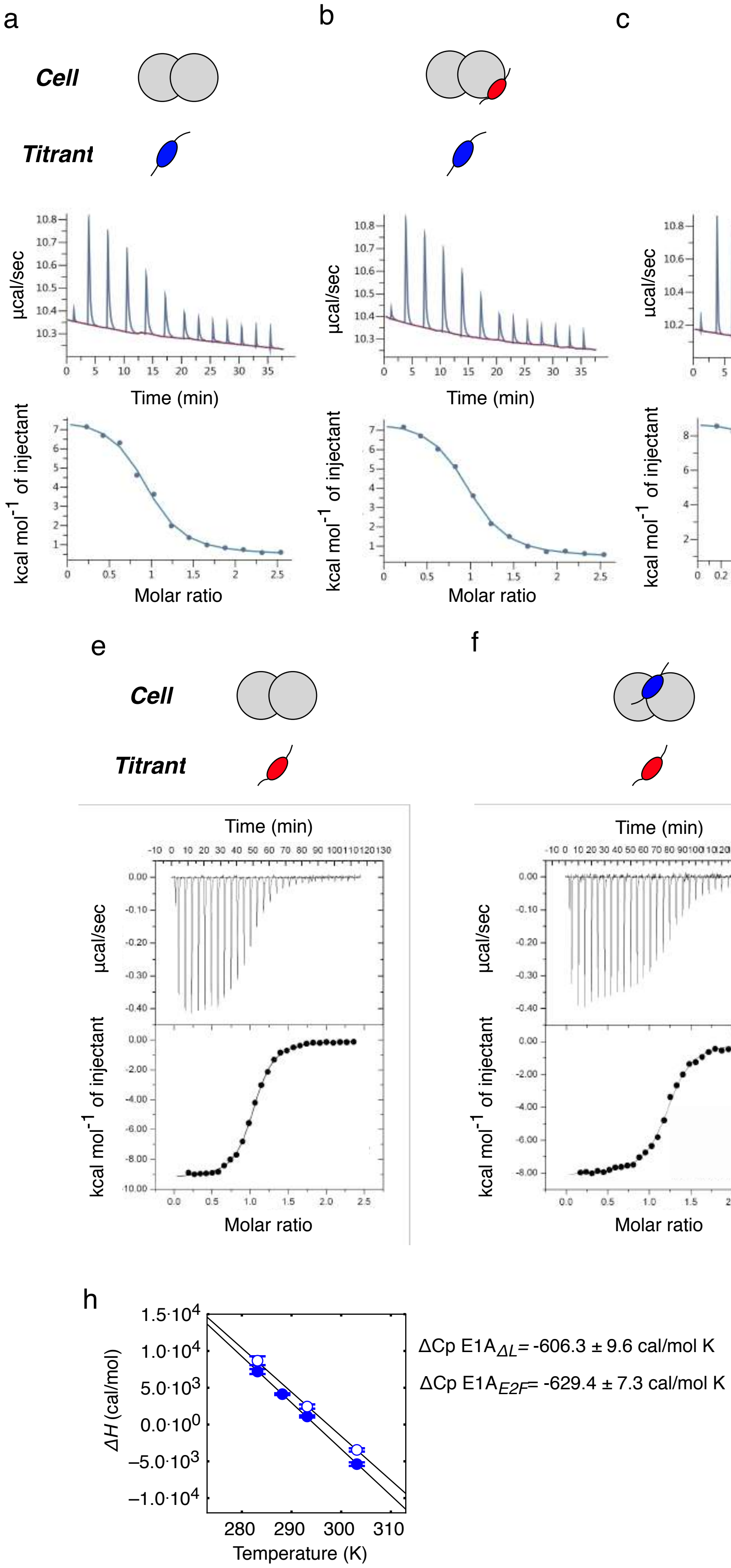

a
b
c
d
Cell
Titrant
μcal/sec
Time (min)
kcal mol-1 of injectant
Molar ratio
e
f
g
Cell
Titrant
Time (min)
μcal/sec
kcal mol-1 of injectant
Molar ratio
h
ΔH (cal/mol)
Temperature (K)
ΔCp E1AΔL= -606.3 ± 9.6 cal/mol K
ΔCp E1AE2F= -629.4 ± 7.3 cal/mol K

**EXTENDED DATA FIGURE 5: Analysis of allosteric effects in the formation of the Rb-E1A complex.** Representative ITC titrations designed to test allosteric effects between the E2F and LxCxE binding sites in Rb. Measurements were performed by loading the cell with Rb or with a pre-assembled complex of Rb with different peptide/proteins containing one of the interacting motifs at stoichiometric concentrations, and titrating with peptide/proteins containing the complementary motif loaded into the syringe. Panels show heat exchanged as a function of time, (upper panel) and the enthalpy per mole of injectant plotted as a function of [peptide or protein]/[Rb or Rb pre-assembled complex] molar ratio (Lower panel, black circles) along with the corresponding fit using a single site binding model (Lower panel, black lines). Binding traces here represented correspond to: **a)** Rb (30 µM, cell) titrated with E1A$_{E2F}$ (300 µM, syringe) at 10 ºC; **b)** [Rb:E1A$_{LxCxE}$] (30 µM, cell) and E1A$_{E2F}$ (300 µM, syringe) at 10 ºC; **c)** Rb (30 µM, cell)) and E1A$_{\Delta L}$(300 µM, syringe) at 10 ºC; **d)** [Rb:E1A$_{LxCxE}$] (30 µM, cell) and E1A$_{\Delta L}$ (300 µM, syringe) at 10 ºC; **e)** Rb (15 µM, cell) and E1A$_{LxCxE}$ (150 µM, syringe) at 20 ºC; **f)** [Rb: E1A$_{E2F}$] (15µM, cell) and E1A$_{LxCxE}$ (150 µM, syringe) at 20 ºC; **g)** [Rb:E1A$_{\Delta L}$] (15 µM, cell) and E1A$_{LxCxE}$ (150 µM, syringe) at 20 ºC. Thermodynamic parameters derived from the fitting are shown in **Extended Data Table 1.** Titrations where Rb was titrated with peptides/proteins containing the E1A E2F motif (a-d) all showed an endothermic behavior at 10 ºC whereas interactions where Rb was titrated with peptides/proteins containing the E1A LxCxE motif (e-g) all showed an exothermic behavior at 20 ºC. A schematic representation of each titration design is shown above the ITC traces: Rb: grey double circle, E2F motif: blue oval, LxCxE motif: red oval. The E1A linker is depicted as a black line. **h) ITC measurements of E1A$_{E2F}$ and E1A$_{\Delta L}$ at different temperatures.** The heat capacity change ($\Delta Cp$) was calculated from the slope of the plot of $\Delta H$ vs temperature. E1A$_{E2F}$: filled blue bars; E1A$_{\Delta L}$: open blue bars. Thermodynamic parameters are reported in **Extended Data Table 5.**

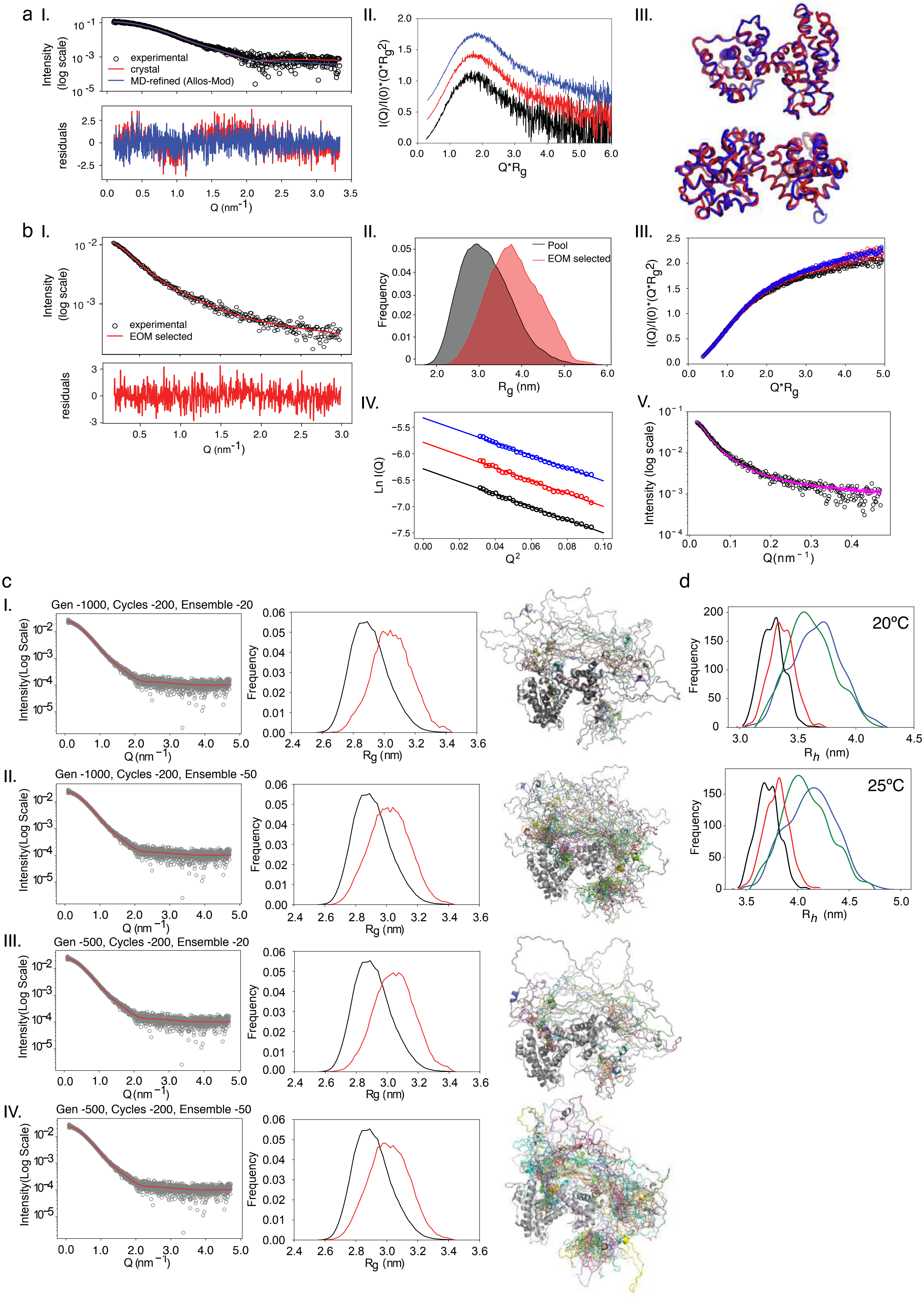

a I.
experimental
crystal
MD-refined (Allos-Mod)
Intensity (log scale)
residuals
Q (nm$^{-1}$)
II.
I(Q)/I(0)*(Q*Rg$^2$)
Q*Rg
III.
b I.
experimental
EOM selected
II.
Pool
EOM selected
Frequency
Rg (nm)
III.
IV.
Ln I(Q)
Q$^2$
V.
c
I.
Gen -1000, Cycles -200, Ensemble -20
Intensity(Log Scale)
II.
Gen -1000, Cycles -200, Ensemble -50
III.
Gen -500, Cycles -200, Ensemble -20
IV.
Gen -500, Cycles -200, Ensemble -50
d
20ºC
25ºC
Rh (nm)

**EXTENDED DATA FIGURE 6: Experimental SAXS intensity profiles of Rb, E1A and of [Rb:E1A] complex fitted using the Ensemble Optimization Method (EOM).**

**a)** I. Comparison of the experimental SAXS intensity profile (black empty circles) with the theoretical SAXS profile obtained from the crystal structure of the unliganded Rb (RbAB domain) (PDB ID: 3POM) (red line) and from a refined model of RbAB where flexible loops were added and the structure refined using the program Allos-Mod-FoXS (blue line) (**See Methods**). Below, residuals of both fits. II. Kratky plots of RbAB measured at 4.0 mg/ml (blue line), 2.0 mg/ml (red line) and 1.0 mg/ml (black line). III. Two orthogonal views of the RbAB domain crystal structure (red) and the optimized RbAB structure with loops modeled (blue) are shown (RMSD = 1.7 Å). **b)** I. SAXS intensity profile of E1A$_{WT}$ (black circles) and the best fit from the EOM method (red line). Below, residual of the fit. II. Rg distribution of the ensemble pool for E1AWT (black area) and the EOM-selected ensemble (red area), where E1A$_{WT}$ samples more extended conformations than these of the pool, which represent a random-coil model. III. Kratky plots of E1A measured at 7.0 mg/ml (blue empty circles), 5.6 mg/ml (red empty circles) and 4.2 mg/ml (black empty circles). IV. Guinier Plots of E1A at the three concentrations tested. V. The SEC-SAXS profile of E1A$_{WT}$ (blue empty circles) and the merged curve from the SAXS experiments at three concentrations (pink line) perfectly overlaid, discarding any aggregation problem. **c)** Theoretical SAXS profiles were computed for a pool of 10250 [Rb:E1A$_{WT}$] complex model structures and compared to experimental SAXS profiles using the EOM method (see Methods). Four fitting conditions are shown using different sub-ensemble pools. From top to bottom: 1000 generations with ensemble size = 20, 1000 generations with ensemble size = 50, 500 generations with ensemble size = 20 and 500 generations with ensemble size = 50. Left panel: experimental SAXS intensity profiles (grey circles) and their fitting using the EOM (red lines). Middle panel: $R_g$ distributions of the pool ensemble (black line) and of each EOM-selected sub-ensemble (red line). Right panel: Representation of the EOM-selected sub-ensembles. The selected sub-ensemble of 1000 generations with ensemble size = 50 describing the experimental SAXS intensity profiles of [Rb:E1A$_{WT}$] is presented in **Figure 3**. **d)** Distribution plots of the $R_h$ calculated for the pool ensemble of 10250 modeled structures of [Rb: E1A$_{WT}$] (black), the EOM-selected [Rb:E1A$_{WT}$] sub-ensemble (red), [Rb:E1A$_{\Delta E}$] (green) and [Rb:E1A$_{\Delta L}$] (blue). Upper panel: Calculations performed at 20 ºC. Lower panel: Calculations performed at 25 ºC.

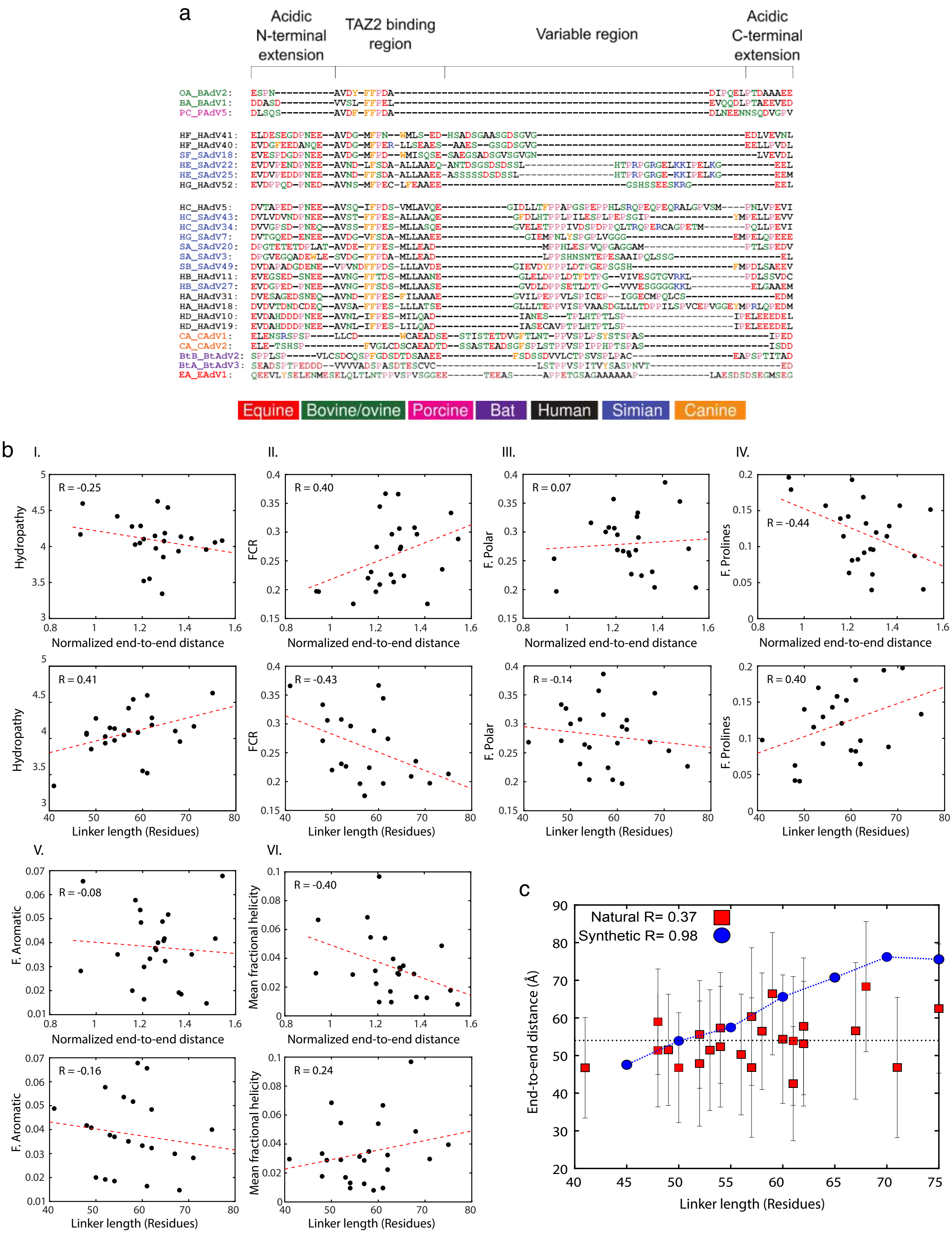

a
Acidic N-terminal extension
TAZ2 binding region
Variable region
Acidic C-terminal extension
Equine
Bovine/ovine
Porcine
Bat
Human
Simian
Canine
b
I.
II.
III.
IV.
V.
VI.
Hydropathy
FCR
F. Polar
F. Prolines
F. Aromatic
Mean fractional helicity
Normalized end-to-end distance
Linker length (Residues)
R = -0.25
R = 0.40
R = 0.07
R = -0.44
R = 0.41
R = -0.43
R = -0.14
R = 0.40
R = -0.08
R = -0.40
R = -0.16
R = 0.24
c
Natural R= 0.37
Synthetic R= 0.98
End-to-end distance (Å)
Linker length (Residues)

**EXTENDED DATA FIGURE 7: Correlation of E1A linker dimensions with sequence-encoded features**
**a)** Global alignment of selected E1A linker sequences generated using a combination of standard sequence alignment tools with an emphasis on conservation of hydrophobic motifs. Amino acids are colored as follows: acidic residues (red), basic residues (blue), polar residues (green), hydrophobic residues (black), aromatic residues (orange) and proline (pink). Colors indicate the hosts of Mastadenoviruses whose E1A sequences were included in the set. From the alignment: the N- and C-terminal extensions of the linkers are highly acidic and conserved. The TAZ2 binding region contains a highly conserved aromatic stretch and a hydrophobic region absent in the shorter bovine and porcine E1A linkers. Sequence length is titrated within the Variable region with polar (G/S/T/N/H/Q) hydrophobic (A/V/I/L/M) and proline residues. **b)** Correlations between distinct sequence parameters and normalized end-to-end distance (upper panels) or linker length (lower panels): Hydropathy (I), fraction of charged residues (FCR) (II), fraction of polar residues (III), fraction of prolines (IV), fraction of aromatic residues (V) and Mean fractional helicity (VI). Most correlations are < 0.3 with several exceptions. The strongest correlation observed is between normalized end-to-end distance or linker length and the net-charge per residue (NCPR) (**Figure 4**). The correlation between fraction of charged residues (FCR) and linker length reflects the fact that the region used to titrate chain length (Variable region in the linker) is largely depleted of charged residues, such that longer chains have a lower fraction of charged residues. This is an important corollary to the correlation with NCPR, as it demonstrates the neutralization of NCPR is not simply via an increase in the fraction of positive residues, but a relative dilution of charged residues. The negative correlation between normalized end-to-end distance vs. proline content (IV, upper panel) may seem counterintuitive at first blush and implies that less extended chains have a higher proline content. This reflects the fact that the variable region is enriched for proline residues – presumably to prevent local folding – such that as the chain gets longer more proline residues are incorporated. The corollary of the latter explanation suggests that there should be a loss of fractional secondary structure as normalized chain length increases, a result borne out when fractional helicity is compared to normalized end-to-end distance (VI, upper panel). The fraction of helicity is generally between 7% and 1%, suggesting that despite this correlation the effect size is relatively small. As a general conclusion, longer chains become overall more proline rich, hydrophobic and less highly charged, leading to a relative compensation in their absolute dimensions. **c)** Linker length control titration experiment. The linker end-to-end distance as a function of linker length for the synthetic sequences (blue circles) compared to the natural sequences (red squares). The collection of random synthetic sequences of different lengths matched the amino acid composition of the HF_HAdV40 linker (**See Methods**).

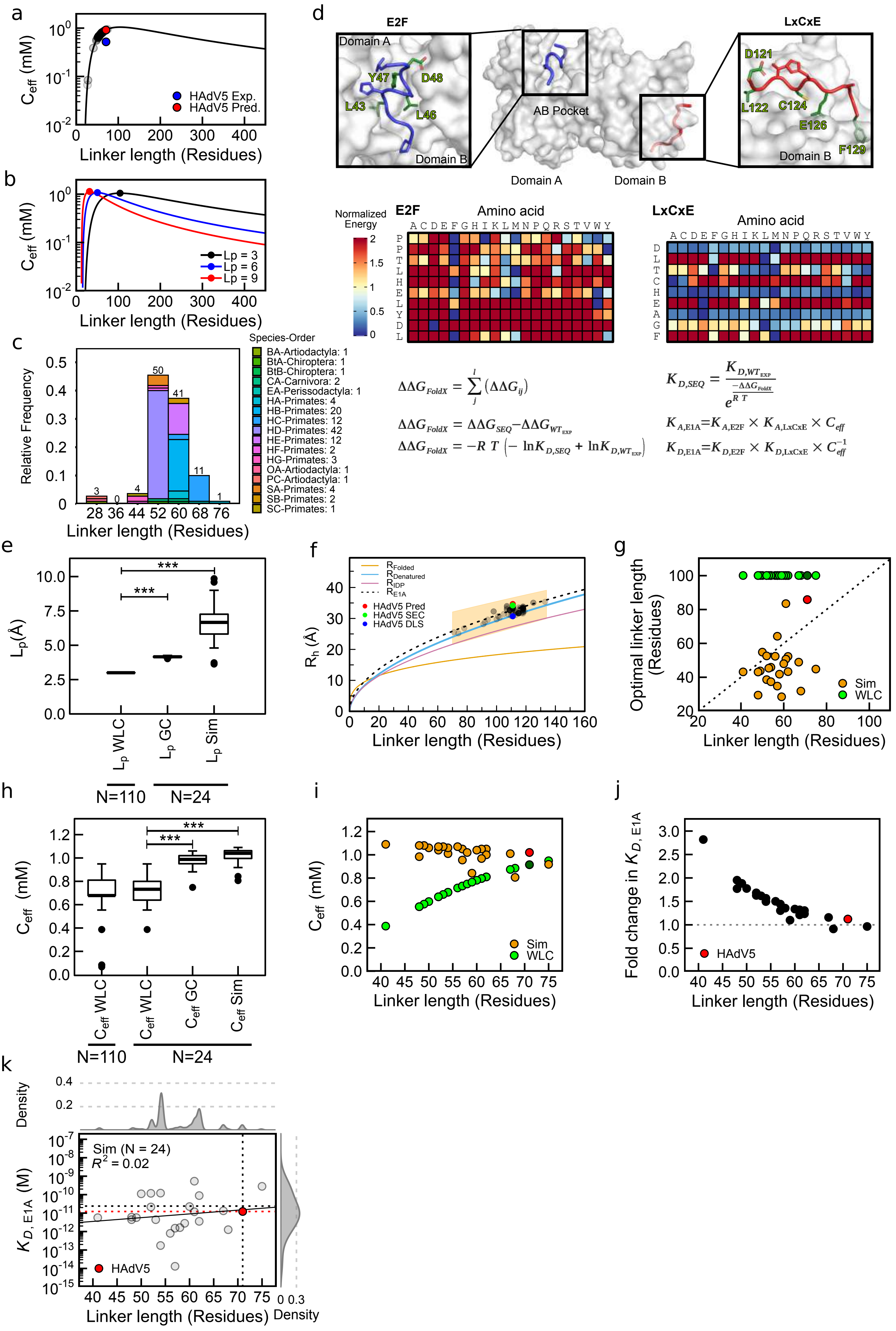

a
Ceff (mM)
HAdV5 Exp.
HAdV5 Pred.
Linker length (Residues)
b
Ceff (mM)
Lp = 3
Lp = 6
Lp = 9
Linker length (Residues)
c
Species-Order
BA-Artiodactyla: 1
BtA-Chiroptera: 1
BtB-Chiroptera: 1
CA-Carnivora: 2
EA-Perissodactyla: 1
HA-Primates: 4
HB-Primates: 20
HC-Primates: 12
HD-Primates: 42
HE-Primates: 12
HF-Primates: 2
HG-Primates: 3
OA-Artiodactyla: 1
PC-Artiodactyla: 1
SA-Primates: 4
SB-Primates: 2
SC-Primates: 1
Relative Frequency
Linker length (Residues)
d
E2F
LxCxE
Domain A
Domain B
Y47
D48
L43
L46
AB Pocket
D121
L122
C124
E126
F129
Normalized Energy
Amino acid
ACDEFGHIKLMNPQRSTVWY
$\Delta\Delta G_{FoldX} = \sum_{j}^{l} (\Delta\Delta G_{ij})$
$\Delta\Delta G_{FoldX} = \Delta\Delta G_{SEQ} - \Delta\Delta G_{WT_{EXP}}$
$\Delta\Delta G_{FoldX} = -R\,T\,(-\ln K_{D,SEQ} + \ln K_{D,WT_{EXP}})$
$K_{D,SEQ} = \frac{K_{D,WT_{EXP}}}{e^{\frac{-\Delta\Delta G_{FoldX}}{R\,T}}}$
$K_{A,E1A} = K_{A,E2F} \times K_{A,LxCxE} \times C_{eff}$
$K_{D,E1A} = K_{D,E2F} \times K_{D,LxCxE} \times C_{eff}^{-1}$
e
Lp(Å)
Lp WLC
Lp GC
Lp Sim
N=110
N=24
f
RFolded
RDenatured
RIDP
RE1A
HAdV5 Pred
HAdV5 SEC
HAdV5 DLS
Rh (Å)
Linker length (Residues)
g
Optimal linker length (Residues)
Sim
WLC
Linker length (Residues)
h
Ceff (mM)
Ceff WLC
Ceff WLC
Ceff GC
Ceff Sim
N=110
N=24
i
Ceff (mM)
Sim
WLC
Linker length (Residues)
j
Fold change in KD, E1A
HAdV5
Linker length (Residues)
k
Density
Sim (N = 24)
R2 = 0.02
KD, E1A (M)
HAdV5
Linker length (Residues)
Density

**EXTENDED DATA FIGURE 8: Global prediction of Rb binding affinity for natural E1A proteins. a)** The predicted effective concentration ($C_{eff}$) for 110 E1A linker sequences was calculated using a worm like chain (WLC) model that considers the linker as an entropic chain and a distance between sites of 49 Å (**See Methods**). The predicted $C_{eff}$ values were grouped near the maximum of the $C_{eff}$ function, showing that E1A linker length might be optimized. **b)** $C_{eff}$ curve using a distance between sites of 49 Å and increasing $L_p$ values. As the $L_p$ value increases, the optimal linker length becomes smaller. **c)** Frequency of linker length for the 110 Mastadenovirus E1A linker sequences analyzed in this work. Lengths are grouped with a bin size of 8 residues. Lower panel: color code used to group hosts Species-Order. The average length is 57 ± 8 residues. **d)** Upper panel: Representation of the E1A protein E2F (blue) and LxCxE (red) motifs bound to the RbAB domain. Insets show details of each motif. Core E2F motif residues L43, L46, Y47 and D48 and core LxCxE motif residues D121, L122, C124, E126 and F129 are depicted as green sticks. The non-core E2F motif residues P40, H41 and L49 (blue sticks), and the LxCxE motif residues T123, H125 and A127 (red sticks) and the A and B Rb subdomains are indicated for reference. Lower panel: FoldX energy matrices of the E2F and LxCxE motifs built from individual complex structures. The values in the matrix were normalized for color code representation in the range 0-2 kcal/mol. Lowest values (blue squares) correspond to mutations that have small destabilizing effects while highest values (red squares) correspond to the most destabilizing mutations. The FoldX matrices were used to calculate a $\Delta\Delta G_{FoldX}$ for each motif and the binding affinity of each sequence ($K_{D,\ SEQ}$) was calculated as explained in Materials and methods. RT was 0.582 kcal/mol. **e)** Distribution of $L_p$ values for the different datasets. The $L_p$ values from all atom simulations ($L_p$ SIM) are more variable and on average larger than the "standard" WLC parameters ($L_p$ WLC) and the Gaussian Chain ($L_p$ GC) simulation (*** $p$-value < 0.001). The whiskers were calculated considering 1.4 IQR. Black dots represent outliers. **f)** Prediction of $R_h$ for 110 natural E1A Motif-Linker-Motif arrangements as a function of linker length (black dots). The predicted $R_h$ was calculated from [4] as $R_h$ = (A * $P_{pro}$+ B)(C * |Q| + D ) * $S_{his}$ * $R_0$ * $N^v$ where A = 1.24, B = 0.904, C = 0.00759, D = 0.963 , $R_0$ = 2.49, v = 0.509. $S_{his}$ = 1 since no polyhistidine tag is present, $P_{pro}$ and |Q| are the observed fraction of proline residues and absolute net charge in the sequence, respectively. The predicted $R_h$ for E1A HAdV5 (red dot) agrees with the SEC (green dot) and DLS (blue dot) experimental measurements. The $R_h$ depends on the number of residues as: $R_h = R_0 * N^v$, where $R_0$ and v are 4.92 and 0.285 for folded proteins ($R_{Folded}$, orange line), 2.33 and 0.549 for chemically denatured proteins ($R_{Denatured}$, blue line) and 2.49 and 0.509 for IDPs ($R_{IDP}$, pink line) [4]. The E1A sequences are predicted to be highly extended as a whole ($R_{E1A}$, black dotted line) according to the average observed $P_{pro}$ and |Q| (0.1 and 26 respectively). **g)** For the data set of 24 linkers, the optimal linker length (peak of the Ceff curve shown in b) is plotted against the actual linker length using $L_p$ = 3 Å ($L_p$ WLC, green dots) or $L_p$ from simulations ($L_p$ SIM, orange dots). For $L_p$ = 3 Å the optimal linker length is fixed (104 residues) and natural linker lengths are predicted to be lower than the optimal linker length (points above the diagonal). However, when $L_p$ SIM is used, the natural linker length is similar or slightly longer than the optimal linker length. HAdV5-E1A values are indicated with dark green and red dots for $L_p$ = 3 Å (WLC) and $L_p$ SIM respectively. **h)** Distribution of $C_{eff}$ values (i) for the different datasets. $C_{eff}$ values using the WLC model are lower and have a wider distribution than those calculated with the $L_p$ predicted from GC and all atom simulations (*** $p$-value < 0.001). The whiskers were calculated considering 1.4 IQR. Black dots represent outliers. **i)** $C_{eff}$ values as a function of linker length for the subset of 24 sequences calculated using the WLC model ($L_p$ = 3 Å) (green dots), or the $L_p$ values predicted from all atom simulations ($L_p$ SIM, orange dots). HAdV5-E1A values are indicated with dark green and red dots respectively. $C_{eff}$ values calculated using $L_p$ SIM are less variable than the $C_{eff}$ calculated with $L_p$ = 3 Å and do not change with linker length. **j)** Fold change in $K_{D,E1A}$ calculated using $L_p$ = 3 Å or $L_p$ SIM ($K_{D,E1A}$ ($L_p$ = 3 Å) / $K_{D,E1A}$ ($L_p$ Sim)) as a function of linker length. Shorter linkers show up to a 2.8-fold change in $K_{D,E1A}$. HAdV5-E1A is indicated with a red dot. **k)** Global Rb binding affinity ($K_{D,E1A}$) as a function of linker length for the dataset of 24 sequences used in simulations ($L_p$ = 3 Å) using the WLC model with the $L_p$ values predicted from the simulations ($L_p$ SIM) (g). Values are in (**Source Data File 2)**. $K_{D,E1A}$ was calculated as explained in Materials and Methods as $K_{D,E1A} = K_{D,E2F} \cdot K_{D,LxCxE} \cdot C_{eff}^{-1}$. The low $R^2$ value indicates that $K_{D,E1A}$ is uncorrelated to linker length. Upper panel: density plot of linker length for 107 E1A sequences. Three short linkers were excluded. Right panel: density plot of $K_{D,E1A}$. Red dot and line: Predicted $K_{D,E1A}$ for HAdV5. The cross lines indicate the linker length and experimentally measured binding affinity of HAdV5-E1A.

References:
1. Uversky VN. What does it mean to be natively unfolded? *Eur J Biochem*. 2002;269(1):2-12. http://www.ncbi.nlm.nih.gov/pubmed/11784292
2. Sherry KP, Das RK, Pappu R V, Barrick D. Control of transcriptional activity by design of charge patterning in the intrinsically disordered RAM region of the Notch receptor. *Proc Natl Acad Sci U S A*. 2017;114(44):E9243-E9252. doi:10.1073/pnas.1706083114
3. Hofmann H, Soranno A, Borgia A, Gast K, Nettels D, Schuler B. Polymer scaling laws of unfolded and intrinsically disordered proteins quantified with single-molecule spectroscopy. *Proc Natl Acad Sci U S A*. 2012;109(40):16155-16160. doi:10.1073/pnas.1207719109
4. Marsh JA, Forman-Kay JD. Sequence determinants of compaction in intrinsically disordered proteins. *Biophys J*. 2010;98(10):2383-2390. doi:10.1016/j.bpj.2010.02.006
5. Perozzo R, Folkers G, Scapozza L. Thermodynamics of protein-ligand interactions: history, presence, and future aspects. *J Recept Signal Transduct Res*. 2004;24(1-2):1-52. http://www.ncbi.nlm.nih.gov/pubmed/15344878
6. Liu X, Marmorstein R. Structure of the retinoblastoma protein bound to adenovirus E1A reveals the molecular basis for viral oncoprotein inactivation of a tumor suppressor. *Genes Dev*. 2007;21(21):2711-2716. doi:10.1101/gad.1590607
