## EXTENDED AND SUPPLEMENTAL DATA for "Conformational buffering underlies functional selection in intrinsically disordered protein regions": 03 SI_TEXT_FOUTEL.pdf

### Supplementary Text Section 1

Our binding results confirm the contribution of flanking regions and phosphorylation to binding of the E1A$_{LxCxE}$ SLiM, as observed in the Human papillomavirus 16 (HPV16) E7 protein [1,2]. The E1A$_{LxCxE}$ SLiM had an affinity similar to E1A$_{E2F}$ ($K_D$ = 105 nM), which increased incrementally upon including the acidic stretch following the motif ($K_D$ = 73 nM), and upon Ser132 phosphorylation ($K_D$ = 20 nM) **(Extended Data Table 1)** leading to an overall 5-fold increase in binding affinity. This range was very similar to that already observed in HPV16 E7 [1,2]. The E1A$_{\Delta L}$ and E1A$_{\Delta E}$ constructs had similar narrow chemical shift dispersion and disorder (**Extended Data Fig.4**) and confirmed that each motif was able to bind independently to Rb (**Fig. 2b III**). However, the high $I/I_0$ values in the region surrounding the mutated motif (average $I/I_0 \sim 0.7$) indicated that the flanking residues only associated to Rb when the core motif was present (**Fig. 2b III**) indicating these regions support binding of the core motif but do not constitute independent binding sites.

### *Supplementary Text Section 2*

To support the binding results, we assessed changes in accessible surface area (*ΔASA)* upon binding using ITC (**Extended Data Fig. 5h and Extended Data Table 4**). We observed no significant increase in the *ΔASA* of E1A$_{\Delta L}$ when compared to E1A$_{E2F}$, indicating the linker did not occlude additional Rb surface from the solvent (**Fig. 2d**). We conclude that stable Rb-linker interactions cannot explain the reduced intensity of the NMR peaks. A plausible explanation may be changes in the conformational exchange rate of the TAZ2 motif due to the formation of transient secondary structure. This possibility is supported by the higher helical content (**Fig. 2b I**), lower disorder propensity (**Fig. 2b IV**) and conservation (**Fig. 2b V**) of this region, which forms a helix upon binding to TAZ2 [3].

### *Supplementary Text Section 3*

The SEC results showed that [Rb:E1A$_{\Delta L}$] and [Rb:E1A$_{\Delta E}$] were more extended than [Rb:E1A$_{WT}$], as expected from a single motif being bound to Rb (**Fig. 3f**). The experimentally measured $R_h$ values were lower than those obtained from computed ensembles (**Fig. 3g, Extended Data Table 3**), suggesting that the linker sampled slightly more compact conformations in these mutants. On the other hand, the $R_g/R_h$ ratio provides a measure of the compactness of a protein [4]. Ratio values of 0.75, 1.0 and 1.50 are expected for compact globules, Flory random chains (FRC) and for excluded volume (EV) chains respectively. Combined SAXS and SEC-SLS measurements confirmed that while Rb had a compact conformation ($R_g/R_h$ = 0.82), *E1A$_{WT}$* was highly extended ($R_g/R_h$ = 1.39). In contrast [Rb:E1A$_{WT}$] had an $R_g/R_h$ = 0.93, consistent with compaction upon bivalent tethering.

#### REFERENCES:

1. Chemes LB, Sanchez IE, Smal C, de Prat-Gay G. Targeting mechanism of the retinoblastoma tumor suppressor by a prototypical viral oncoprotein. Structural modularity, intrinsic disorder and phosphorylation of human papillomavirus E7. *FEBS J.* 2010;277(4):973-988. doi:10.1111/j.1742-4658.2009.07540.x
2. Palopoli N, Gonzalez Foutel NS, Gibson TJ, Chemes LB. Short linear motif core and flanking regions modulate retinoblastoma protein binding affinity and specificity. *Protein*

*Eng Des Sel*. 2018;31(3):69-77. doi:10.1093/protein/gzx068
3. Ferreon JC, Martinez-Yamout MA, Dyson HJ, Wright PE. Structural basis for subversion of cellular control mechanisms by the adenoviral E1A oncoprotein. *Proc Natl Acad Sci U S A*. 2009;106(32):13260-13265. doi:10.1073/pnas.0906770106
4. Sherry KP, Das RK, Pappu R V, Barrick D. Control of transcriptional activity by design of charge patterning in the intrinsically disordered RAM region of the Notch receptor. *Proc Natl Acad Sci U S A*. 2017;114(44):E9243-E9252. doi:10.1073/pnas.1706083114
