## EXTENDED AND SUPPLEMENTAL DATA for "Conformational buffering underlies functional selection in intrinsically disordered protein regions": 08 SOURCE DATA_FOUTEL_Description.pdf

**SOURCE DATA 1. Table of All-Atom Simulation parameters.** For 27 selected E1A sequences representative of *Mastadenovirus* phylogeny, a series of structural parameters were derived from all-atom simulations (Columns D-G) and other sequence parameters were calculated using the local CIDER software package [ ] (Columns H-Y). The species and type names for each E1A sequence is indicated as the species abbreviation followed by the type abbreviation. Three out of 27 sequences were discarded due to their short sequence length (< 30 residues) and the others were selected to perform statistical analysis whose conclusions were used throughout this work.

**SOURCE DATA 2. Table of E1A sequences and predicted Ceff and binding affinities.** ***Sheet 1 Sequences***: For 116 E1A sequences representative of *Mastadenovirus* phylogeny, we identified the E2F and LxCxE motifs. Both motifs were present in 110 sequences and the linkers were defined between the first position after the end of the E2F motif to the last position before the LxCxE motif. Predicted binding affinities using the FoldX matrix energy, the WLC model with and without standard parameters and from all-atom and gaussian chain simulations are presented. ***Sheet 2 KDs Statistics:*** Permutation tests performed to test for significant differences between binding affinities. ***Sheet 3 KDs CI 99%***: 99% confidence intervals calculated using bootstrapping to test for differences with the E2F2 binding affinity. ***Sheet 4 Ceff statistics and Sheet 5 Lp statistics:*** Permutations tests performed to test for significant differences between both $C_{eff}$ datasets and $L_p$ datasets. ***Sheet 6 E2F FoldX Matrix and Sheet 7 LxCxE FoldX Matrix:*** Matrices used for calculating binding affinities.

**SOURCE DATA 3. Model of E1A$_{LxCxE}$ bound to Rb.** PDB data file of a model of the E1A$_{LxCxE}$ motif in complex with Rb built using FlexPepDock and the structure of the HPV E7 LxCxE motif bound to Rb (PDB: 1GUX).

**SOURCE DATA 4. E1A Alignment.** Alignment of 116 E1A sequences including the region encompassing the E2F motif, linker region and LxCxE motif.
